# Digital nanoreactors for control over absolute stoichiometry and spatiotemporal behavior of receptors within lipid bilayers

**DOI:** 10.1101/2022.10.04.509789

**Authors:** Vishal Maingi, Zhao Zhang, Chris Thachuk, Namita Sarraf, Edwin R. Chapman, Paul W.K. Rothemund

## Abstract

Interactions between membrane proteins are essential for cell survival and proper function, but the structural and mechanistic details of these interactions are often poorly understood. Even the biologically functional ratio of protein components within a multi-subunit membrane complex—the native stoichiometry—is difficult to establish. We have demonstrated digital nanoreactors that can control interactions between lipid-bound molecular receptors along three key dimensions: stoichiometric, spatial, and temporal. Each nanoreactor is based on a DNA origami ring, which both templates the synthesis of a liposome and provides tethering sites for DNA-based receptors. Receptors are released into the liposomal membrane using strand displacement and a DNA logic gate measures receptor heterodimer formation. High-efficiency tethering of receptors enables the kinetics of receptors in 1:1 and 2:2 absolute stoichiometries to be observed by bulk fluorescence in a plate reader which in principle is generalizable to any ratio. Similar ‘single molecule in bulk’ experiments using DNA-linked membrane proteins could determine native stoichiometry and the kinetics of membrane protein interactions for applications ranging from signalling research to drug discovery.

## Introduction

Many cellular functions are mediated by signalling events triggered by protein-protein encounters occurring within lipid bilayer membranes.^1^ Understanding membrane protein interactions and their downstream effects often provides direct and important insight into how cells function on the molecular level. Membrane protein interactions trigger countless cascades of events essential to cellular function, yet for many membrane proteins we lack even a basic understanding of what structural arrangement is necessary to trigger these events. However, it is often difficult to establish whether the active form of an integral membrane protein is a monomer or oligomer (a complex containing two or more interacting partners), or which of many potential homomeric or heteromeric complexes is physiologically relevant.^2^ Basic characterization of the biologically active oligomeric state of membrane proteins is a prerequisite to understanding their function^3–5^ and is useful for drug discovery,^6,7^ dissecting the molecular mechanism of pathogenic processes,^8^ and elucidating the role of transient membrane protein interactions.^9^

Existing experimental approaches for characterization of the oligomeric state each have their limitations: polyacrylamide gel electrophoresis cannot replicate the native lipid environment and can itself introduce artifactual dimers;^10^ chemical cross-linking can be employed to stabilize oligomers under non-native conditions at the risk of introducing artifactual dimers from nonspecific reactions;^11^ bulk Förster resonance energy transfer (FRET) data is concentration sensitive, and must be carefully corrected to account for potential FRET between oligomers,^12^ single molecule fluorescence photobleaching and FRET methods can exquisitely resolve features of oligomers but are technically challenging,^11–16^ and mass spectrometry requires detergents for sample preparation and expensive instrumentation.^17^ Often, to definitively characterize the oligomeric state, it is necessary to combine multiple analytical approaches, adding time and complexity. An ideal experimental platform for membrane protein interactions would avoid the drawbacks of the methods above, enable the study of isolated proteins in a cell-free yet native lipid environment, and measure real time kinetics and dynamics of their interactions. Further, the platform would simultaneously provide precise *stoichiometric, spatial*, and *temporal* (S^2^T) control: exact numbers of monomeric proteins would begin in a well-separated initial configuration within a well-defined reaction volume, and their triggered release could be used to time the beginning of the experiment.

One path to such an ideal platform is DNA nanotechnology, which has recently been used to construct a number of “custom instruments for biology”^18–25^ wherein DNA nanostructures are designed from the beginning to ask exactly the experimental question at hand. The construction of custom molecular instruments has been enabled by the versatility of DNA nanotechnology: DNA can be folded^26^ or assembled into 2D^27,28^ or 3D^29,30^ shapes, these shapes can be programmed to create reconfigurable devices and machines^31^, and can be decorated with a variety of functional groups, *e.g*. proteins^32^ and polymers,^33^ whose position can be controlled in 0.34 nm steps. This has enabled S^2^T control in the context of surface chemical reaction networks on DNA origami^34–38^, where reactants hop from one periodic lattice site to the next. Critical to extending S^2^T control to fluid bilayers are commercially available and custom-made hydrophobic modifications to that interface DNA with lipid membranes: they have been used by many research groups to engineer and study DNA-lipid systems^39–41^ with applications varying from artificial nanopores,^42,43^ to membrane sculpturing,^44^ nanodiscs,^45–47^, DNA circuits,^48–51^, control of liposome fusion, and artificial cells. Yet so far, no such system has achieved full S^2^T control on a lipid bilayer.

Here, our approach is to use DNA nanotechnology to build a hybrid DNA-lipid instrument, a **D**NA **O**rigami-templated **L**iposome (DOL)^39^, which provides a generic assay platform to orchestrate and measure the interactions between reacting species in a single lipid bilayer. To validate our platform, we used membrane-anchored DNA complexes, which we term DNA receptors, as models for membrane proteins. We exploited several strategies to create the first membrane-based platform that achieves full S^2^T control. First, we used a well-defined and addressable structure of DNA origami^27^ nanocage^59^ to exert absolute stoichiometric^60–62^ and spatial control over the DNA receptors. Building on previous work^39,59^ that demonstrates that guest liposomes of well-defined size can be templated within DNA origami cages, we arranged discrete numbers (e.g. two or four) of DNA receptors at precise distances (e.g. 45 nm) along the circumference of a liposome-filled cage to create a well-defined initial state (Figure 1A and B, Step 1). Cholesterol on the DNA receptors bound them to the liposome, and tethers between the receptors and the cage served to protect them and keep them from reacting until desired. Next we used toehold-mediated strand displacement (TMSD^63–65^) to both provide temporal control via triggered reaction initiation (Figure 1A and B, Step 2) and to create a DNA logic gate^63^ that outputs a fluorescent signal to measure the extent of DNA receptor heterodimerization (Figure 1A and B, step Step 3). Because the logic gate requires simultaneous interaction of both receptors with a reporter complex, our system models a ligand-induced protein dimerization process. Measurement of DNA receptor interaction kinetics for two different absolute stoichiometries, both on the DOL and in solution, show that: (1) we achieved digital control over the number of receptor complexes localized to the DOL, (2) receptors interacted primarily within a single DOL rather than between DOL, and (3) DOL-bound receptors reacted with an effective rate constant that is 2800-fold higher than that measured in solution. Thus, DOL can be thought of as digital nanoreactors—defining, isolating, and concentrating reactions between membrane-bound receptors.

**Figure 1:**
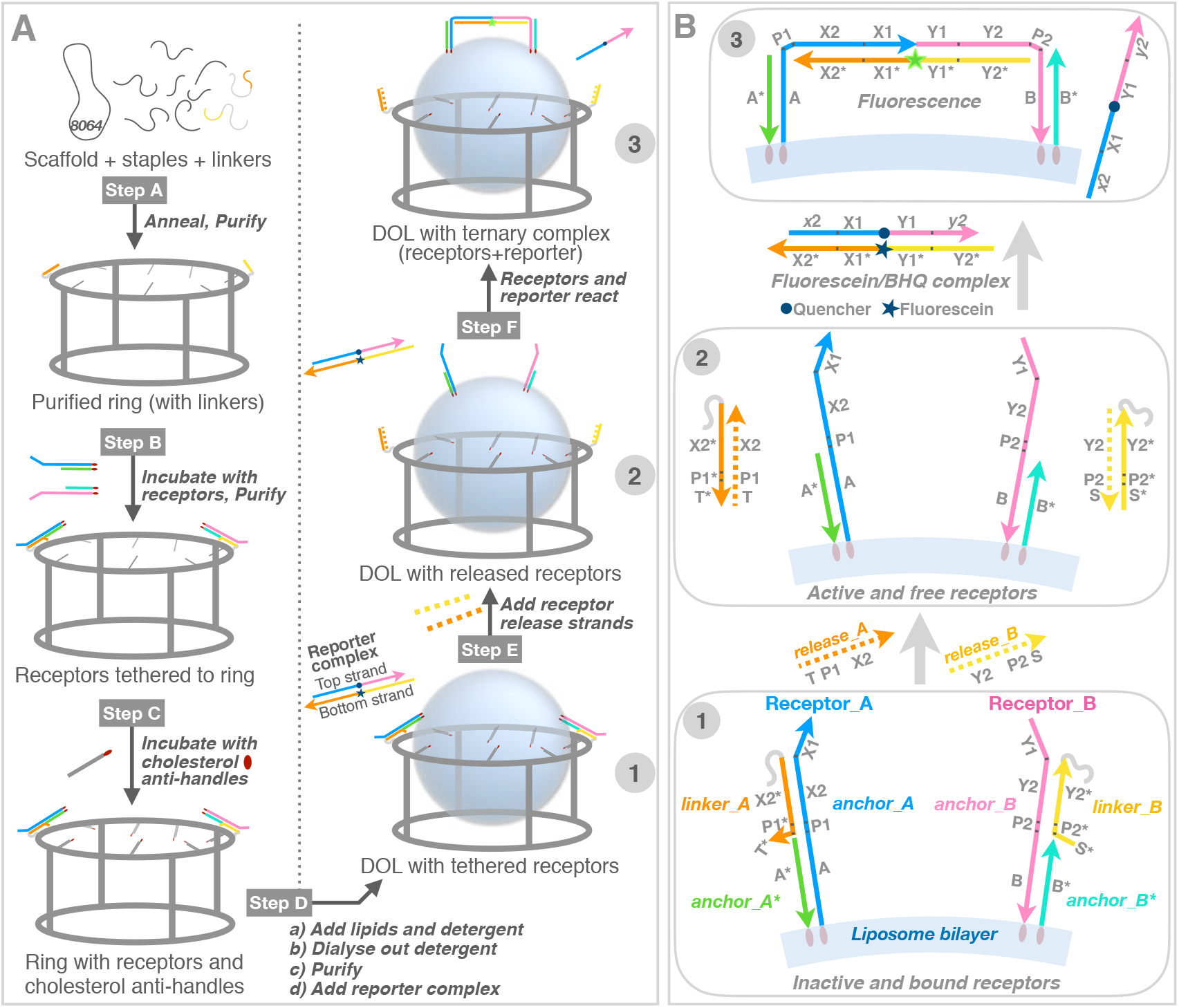
DOL synthesis and DNA circuit logic. **(A)** DOL assembly (left column) and DNA receptor interaction (right column). Step A: DNA scaffold (8064 nucleotides; grey loop), regular staple strands (grey segments), and linker-extended staples (having orange and yellow sections) were annealed; excess staples were removed. Step B: Two types of DNA receptors modified with cholesterol (red ovals) were tethered to the ring and rings were repurified. Step C: Rings were incubated with cholesterol-modified antihandles (grey lines with red ovals) Step D: Lipids and detergent were added; subsequent dialysis removed detergent and seeded liposome formation (blue spheres) on rings to create DOL. **(B)** Stepwise operation of a DNA circuit for the receptor release and interaction measurement. Step labels 1-3 correspond to labels in the right column of **A**. A zoomed segment of the liposome bilayer is shown. Initially (Step 1) both receptors are inactive and bound to the ring (not shown) via *linker_A* and *linker_B* (themselves attached to the ring via a short section of gray polyT). The inter-receptor distance (~45 nm) is not shown to scale. Receptors were detethered (Step 2) by adding release strands complementary to the linkers; domains T* and S* provided toeholds for this reaction. Released receptors diffuse freely within the bilayer but do not interact. Receptor interaction (Step 3) is mediated by a reporter complex consisting of a top strand with internal quencher (dark blue circle) and a bottom strand with an internal fluorophore (star; dark blue when quenched or green when fluorescent). Table 1 gives domains and sequences for all circuit components; Supplementary Table S1 gives each domain’s role).

## Synthesis and Circuit Design

### The DOL platform

Figure 1A summarizes our DOL synthesis strategy (Figure 1A left) and its use for controlling DNA receptor interactions (Figure 1A right, Supplementary Figure S1). A pool of staple strands, including special staples with linker extensions was annealed with a circular DNA scaffold (Step A) to assemble a cage-like DNA origami comprising two interconnected rings; here we refer to this entire structure simply as a ring. The linker-extended staples were designed to specify the number, position, and type of DNA receptors that were attached to the ring in the next step. In particular, the sticky end overhangs presented by each linker determine which receptor type will bind at a particular position on the ring. Two linkers are shown in Figure 1A, suitable for 1:1 receptor absolute stoichiometry; four linkers were used for 2:2 receptor absolute stoichiometry. To remove excess staples and undesired higher-order structures, the reaction products were purified via rate-zonal ultracentrifugation (separating by size, Supplementary Figure S1B). Next, preformed DNA receptors were attached to the rings by an isothermal incubation (Step B); excess receptors were removed in a second rate-zonal ultracentrifugation (Supplementary Figure S1C). Additionally, at least thirty staples on the ring, termed handles, carry extensions designed to bind complementary cholesterol-modified DNA strands termed anti-handles. Anti-handles were attached to rings in a second isothermal incubation (Step C). The cholesterol-modified rings, with tethered receptors, were next mixed with lipids and detergents (Step D). During a follow-up detergent removal process, the cholesterol modifications served as seed for the formation of a liposome on each ring, creating DOL. The resultant mixture, containing undesired free liposomes and DOL, were purified using isopycnic ultracentrifugation (separating by density, Supplementary Figure S1D). Fractions containing fully assembled DOL (see Supplementary Section 1) were used to analyze DNA receptor interactions.

### DNA receptors and their interaction logic

Figure 1B and Table 1 show the domain-level representation of our two different types of DNA receptor complexes (Receptor_A and Receptor_B). We explain domain level details for Receptor_A; Receptor_B has the same domain level structure, but with different sequences. Here and throughout, strand names are italicized, and domain names are bolded. Supplementary Table S1 more extensively describes domains and their roles. Sequence design and analysis were done with NUPACK^66^, which employs SantaLucia nearest-neighbor parameters^67^, assuming 1 M Na^+^ at 25 °C and using default dangle parameters.

**Table 1:**
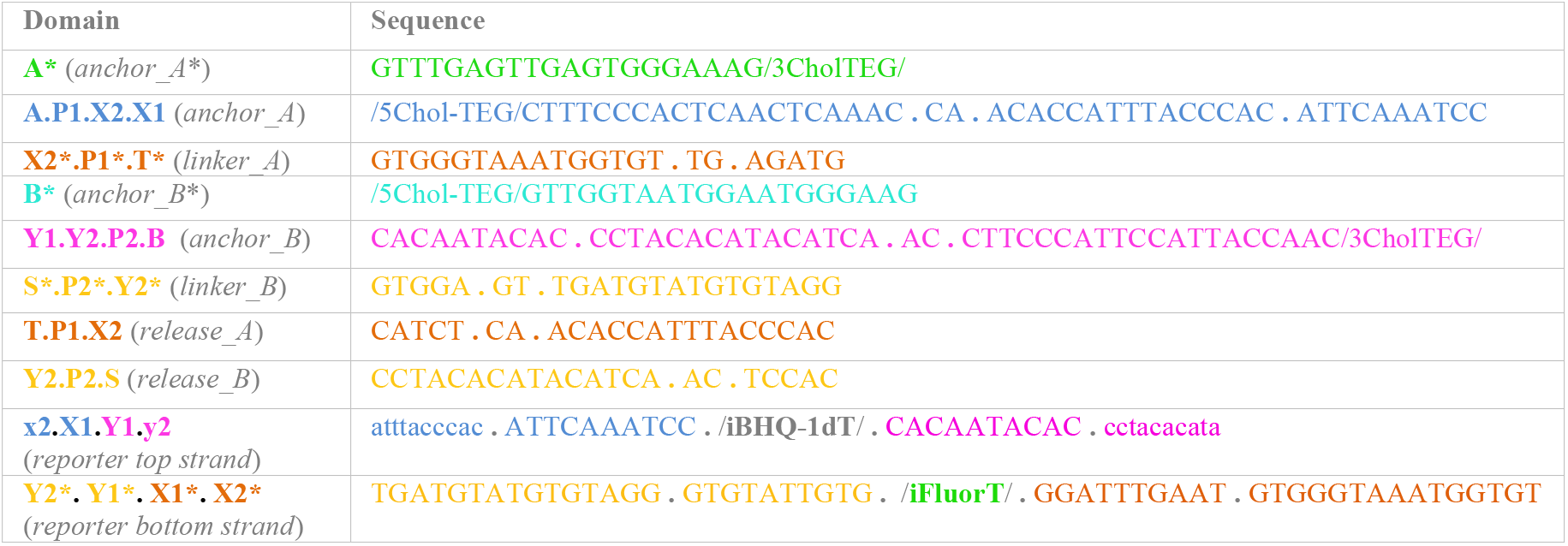
Domain-level sequences (5’—3’) of two different types of DNA receptor and reporter complexes. See Figure 1 for domain-level diagrams of different complexes and Table S1 for more extensive description of domains and their roles. iBHQ-1dT (IDT commercial code) is a black hole quencher and iFluorT (IDT commercial code) is a fluorescein and both conjugated to internal T nucleotides, Chol-TEG is a TEG linked cholesterol molecule (IDT commercial code). Note that domains labeled with lower case are partially complementary to their upper-case counterparts. E.g. **x2** (reporter top strand) is a shortened version of **X2** and is only partially complementary to **X2\*** (reporter bottom strand). Colour codes used here correspond to the same coloured domains shown in Figure 1.

Receptor_A is composed of two cholesterol-modified oligonucleotides, *anchor_A* and *anchor_A*\*. Domain **A** (in *anchor_A*) is complementary to domain **A**\* (the only domain in *anchor_A*\*); together these domains serve the purpose of membrane anchoring via their cholesterol modifications. In general, the use of two cholesterols provides more stable association of DNA complexes with membranes than does a single modification.^49,54^ **P1X2** in *anchor_A* hybridizes with **P1*X2\*** in *linker_A* (an extension from a staple strand in the ring) which tethers the receptor to the ring during DOL assembly (Step B, Figure 1A). After synthesis, in Step 1 (Figure 1B), receptors are separated on the ring by ~ 45 nm.

In Step 2, addition of *release_A* strand, results in TMSD release of Receptor_A, as initiated by the hybridization of domain **T** with toehold **T\*** on *linker_A*. The subsequences **TP1X2** (*release_A*) and **X2*P1*T\*** (*linker_A*) are fully complementary, and thus their full hybridization, after TMSD of **P1X2**, is thermodynamically more favorable and essentially irreversible. The released Receptor_A has a free unpaired subsequence **P1X2X1** and, similarly, after the addition of *release_B*, the Receptor_B has an unpaired **Y1Y2P2** subsequence. By design, **P1X2X1** (NUPACK-calculated free energy ΔG° = −0.27 kcal mol^-1^; shows little predicted secondary structure) and **Y1Y2P2** (NUPACK-calculated free energy ΔG° = 0; unstructured) are not predicted to hybridize (NUPACK reports no bound complex at experimentally relevant concentrations); thus, released Receptor_A and Receptor_B are unlikely to interact with each other.

Released receptors can only interact (Step 3) in the presence of a reporter complex (‘ligand’), which is a fluorophore-quenched duplex comprising a top strand containing an internal quencher (black hole quencher; BHQ) and a bottom strand containing an internal fluorophore (fluorescein). The bottom strand of the reporter (**Y2*Y1*X1*X2**\*) has two five-nucleotide toeholds: **X2\*** initiates binding of Receptor_A to reporter via **X2** in *anchor_A* and **Y2\*** initiates binding of Receptor_B to reporter via **Y2** in *anchor_B*. Overall, the formation of a ternary complex by Receptor_A, Receptor_B, and the reporter’s bottom strand is very similar to the cooperative hybridization reaction reported by Zhang.^68^ Note that intermediate states formed by either receptor individually with the reporter complex (*i.e*. states A^i^ and B^i^ in Supplementary Figure S2) are thermodynamically less favorable than the reactants, and thus sequester very little of either receptor.^68^ Further, formation of intermediate states, which are kinetically reversible, does not result in dequenching of the fluorophore (Figure 4, discussed below). Successful dequenching of the reporter complex (and resulting fluorescence) is only possible when both the receptors are present to cooperatively displace the BHQ-containing top strand. **P1** and **P2** domains of the ternary complex remain unpaired, acting as flexible hinges.

## Results and Discussion

### Intra-DOL receptor interactions

Implementing the DNA logic gate shown in Figure 1B, we explain here interactions between two receptors, one Receptor_A and one Receptor_B per DOL (DOL^1A1B^), initially tethered at distal ends of the ring and anchored in the liposome membrane with their cholesterol ends (Figure 2A, left). To set up a plate reader experiment, the reporter complex (final concentration 4.7 nM) was first mixed with purified DOL^1A1B^ fraction and then the fluorescence intensity was initially measured for ~ 7 h (Figure 2D, blue curve). No increase in fluorescence was observed during this phase because the lipid anchored receptors remain inactive and tethered to the ring via linker strands. Note that linker strands serve the dual purpose of tethering as well as protecting the reactive domains of the receptors. This initial period (7 h) of measurement served as a quality check of our overall purification process. If our purification method of getting rid of untethered reactive receptors was not successful, we would expect to see a rise in signal during this phase. Any unbound and thus active receptors, possibly in solution or on DOL, with their reacting domains **P1X2X1** (in *anchor_A*) and **Y1Y2P2** (in *anchor_B*) can interact with the reporter complex in solution to generate fluorescence. But no significant change in fluorescence was observed, indicating that our purification protocol successfully removed most of the unbound excess receptors (see related discussion in Supplementary Information Section 2).

**Figure 2:**
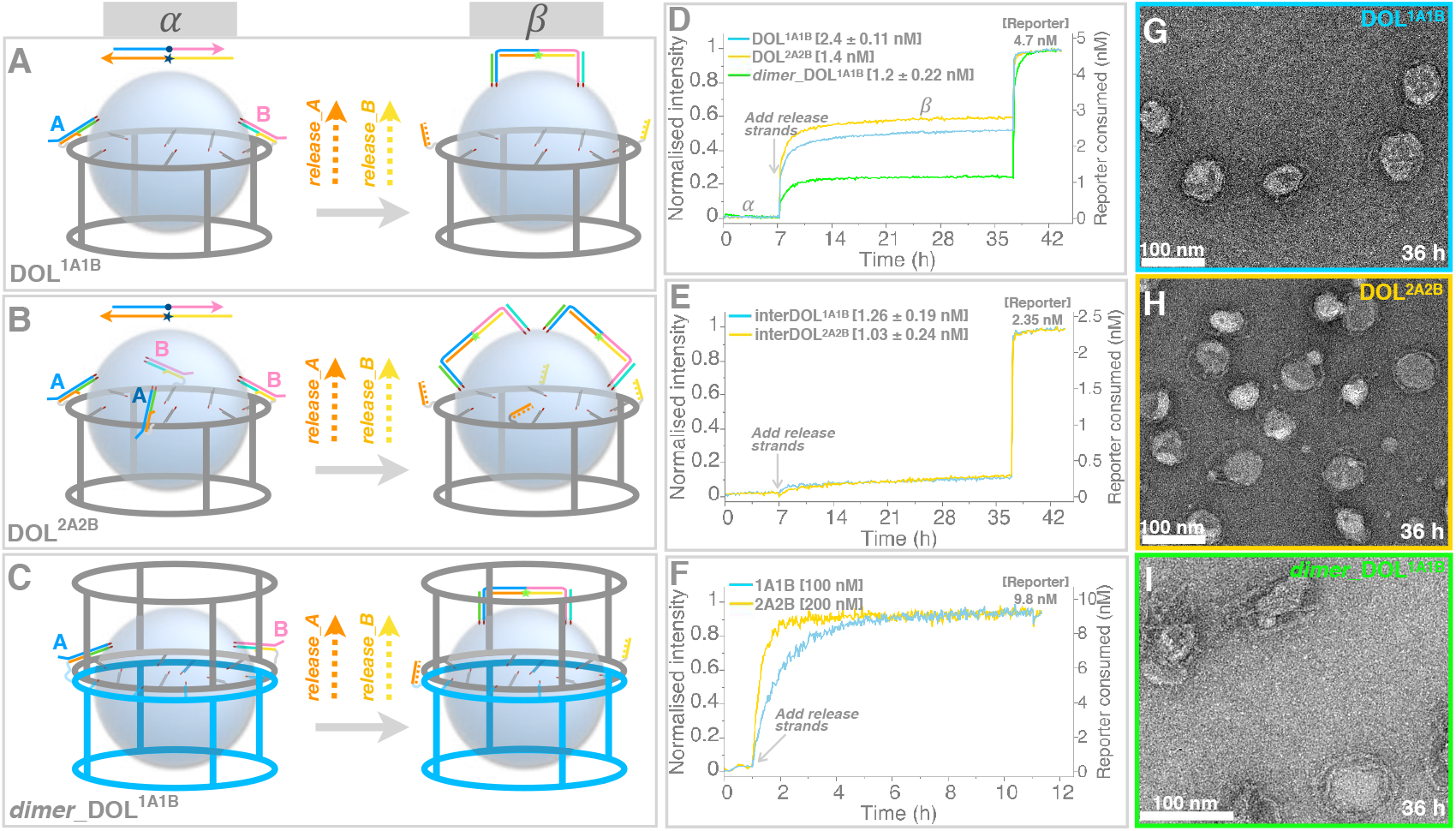
Receptor reactions on three DOL variants. (A-C) Various platforms studied here by varying the number of receptors or rings. In each case also shown (right side of arrow) the number of ternary complexes depending on the initial number of receptors tethered on a DOL platform (left side of arrow). The DNA reporter circuit logic (Figure 1B) remains the same in all the cases. Labels α (left) and β (right) represent the states corresponding to fluorescence intensity curves shown in **D-F**. **(D-F)** Kinetics curves acquired from plate reader experiments shown for receptor interaction event on the same surface of DOL (intra-DOL): DOL^1A1B^ (cyan curve, pooled fraction 3+4, two repeats averaged), DOL^2A2B^ (orange curve, fraction 5, single repeat), and *dimer*_DOL^1A1B^ cases (green curve, fraction 6, two repeats averaged). Initial 7 h has DOL with reporter complex (4.7 nM). After 7 h release strands (100 nM) were added. DOL concentrations are the saturation endpoints, with single standard deviation for two repeats where performed, are given in square brackets. To measure the maximum available fluorescence for purposes of normalization, unreacted reporter was unquenched by adding an excess of stimulant strands at ~ 36 h at 100 nM (*anchor_A* and *anchor_B* without cholesterol modifications) evident as a quick spike in fluorescence. States labelled as α (before 7 h) and β (after 7 h) are shown as cartoon representations **A**, **B**, **C** for each case. **(E)** Kinetics curves shown for receptor interaction between two different DOLs (inter-DOL) each containing only one receptor type: inter-DOL^1A1B^ and inter-DOL^2A2B^. In all cases pooled fraction 3+4 were used and two repeats were performed (averaged curves shown). Plate reader experiment details similar to **D**. Concentrations were estimated from TEM data (see Supplementary Information Section 4). **(F)** Kinetics curves shown for receptor interaction in solution. Receptor complexes were made with linker and anchor strands without cholesterol modifications (Receptor_A consists of *linker_A* and *anchor_A*\* and *anchor_A*, similar for Receptor_B). Plate reader experiment details are similar to **D** and receptors are activated by adding release strands. For 1A1B case each receptor 100 nM, release strands 900 nM and for 2A2B case these were at 200 nM and 1800 nM respectively, reporter complex was at 9.8 nM for both cases. Adding excess of stimulant strands did not show any further spike in fluorescence as all of the reporter molecules were consumed by receptors already in excess. (**G**-**I)** TEM images for the samples taken after completion of plate reader experiment (after ~ 36 h) for the DOL cases in **D**. TEM images for interDOL cases in **E** are shown in Supplementary Figure S8.

After 7 h, a mix of *release_A* and *release_B* (both at 100 nM final concentration) was added, which triggered the release of both the receptors on the surface of lipid bilayer. Through cooperative hybridization, both the active receptors react with the reporter complex to completely displace the BHQ top strand forming one ternary complex per DOL (Figure 2A, right). As a result, a quick rise in fluorescence was observed which almost saturated within ~ 3.5 h of releasing the receptors. Similarly, using the same DOL platform but with two additional linkers, we studied another case where two Receptor_A and two Receptor_B were tethered per DOL which form two ternary complexes per DOL (DOL^2A2B^, Figure 2B). Figure 2D, orange curve, shows the fluorescence kinetics for DOL^2A2B^ case. In all the cases, here and other cases discussed later, to determine whether all the reporter complex has been consumed or not, excess of *anchor_A* and *anchor_B* strands (without cholesterol modifications, and 100 nM final concentration for each) was added and then the fluorescence was measured for another 4 to 6 h. This helped us to normalize the data and also this procedure provides an indirect way to measure DOL concentration by knowing the fraction of reporter consumed (Supplementary Information Section 3). Thus, in all these cases, the fluorescence saturation achieved at *c.a*. 36 h is related to the reporter complex consumed by receptors present on DOL, and is thus dependent on the DOL concentration in a particular fraction used for analyses as explained further.

After the purification step (Step D, Supplementary Figure S1D) the collected fractions are expected to have different concentrations of DOL (with tethered receptors). Thus, after adding *release_A* and *release_B* the maximum fluorescence intensity that can be achieved in each fraction is proportional to the DOL concentration. For example, electron microscope images and fluorescence intensity of DOL^1A1B^ suggested that DOL concentration is higher in fractions 3 and 4 than in fraction 5 as shown in Figure 3 and Supplementary Figure S3 (see related discussion in Supplementary Section 1). So, fractions 3 and 4 were pooled to have more volume for analyses. In Figure 2D, the kinetics curve for DOL^1A1B^ is shown for the pooled fractions 3 and 4, which has higher fluorescence intensity, thus higher DOL concentration, than fraction 5 (compared in Supplementary Figure S4A). On the other hand, in the case of DOL^2A2B^ (Figure 2D) the fluorescence kinetics curve is shown for fraction 5, but the pooled fractions (3+4) consumed the reporter complex completely (Supplementary Figure S4A) which implies that total receptor concentration in pooled fractions (3+4) was at least high as the reporter concentration (see related discussion in Supplementary Information Sections 3 and 4). Thus, the fluorescence curve for combined fraction (3+4) in DOL^2A2B^ case was not used to perform additional analyses (*e.g*. measuring concentration or deriving rate constants).

**Figure 3:**
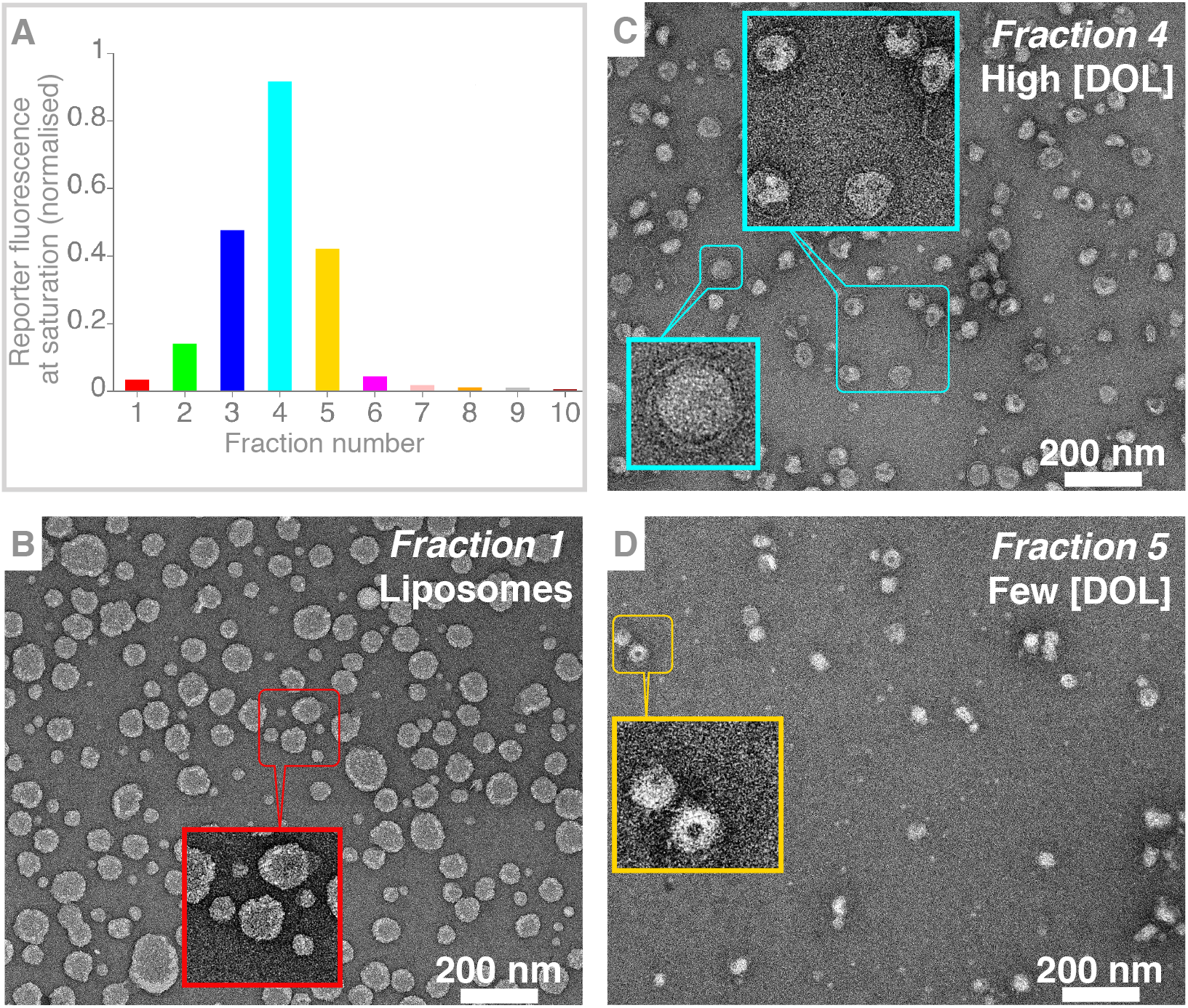
Analyzing different fractions for DOL^1A1B^. **(A)** Different fractions collected after isopycnic DOL purification (see Supplementary Section 1 and Supplementary Figure S1D) were analyzed with plate reader experiment set up similar to Figure 2D. Normalized saturation for each fraction is shown as a bar plot. TEM images for fractions 1, 4 and 5 are shown in **B**, **C**, **D** respectively and remaining fractions are shown in Supplementary Figure S3.

We also explored the situation where two types of receptors were tethered to two rings, which later dimerized and together templated a liposome (Supplementary Information Section 5, Supplementary Figure S6 and Figure S7). Similar to the above cases, both receptors were released on the template liposome bilayer and fluorescence was measured in the presence of reporter complex (Figure 2C and 2D). The dimer platform demonstrates that reactants can come from two different rings, which could be potentially suitable for specific applications (discussed in Conclusions section).

### Inter-DOL receptor interactions

The main purpose of the DOL platform is to control and quantify single-molecule isolated interactions between receptors on the same lipid bilayer surface (intra-DOL) with minimal cross-talk among the DOLs in bulk solution. Thus, it is essential to determine any contribution originating from one receptor interacting with the other on two different DOLs (inter-DOL). We created DOL having only a single type of receptor, which allowed us to study receptor interactions purely as inter-DOL reactions. For example, to evaluate possible inter-DOL interactions in the case of DOL^1A1B^ (intra-DOL) we assembled DOL^1A^ and DOL^1B^ individually, and then mixed (referred as interDOL^1A1B^) equal volumes of their purified fractions (pooled fractions 3+4 each case), and performed similar plate reader measurements as described above for intra-DOL cases. Overall, Figure 2E shows that the inter-DOL reaction rate is slower than the intra-DOL interaction. This implies that most of the fluorescence signal obtained in the intra-DOL case, which has faster reaction kinetics, is due to receptors anchored on the same surface. Similarly, comparing inter-DOL interaction of DOL^2A^ and DOL^2B^ (interDOL^2A2B^) with intra-DOL DOL^2A2B^ faster kinetics was observed in DOL^2A2B^ (Figure 2D and 2E).

### Kinetics

Figure 2D-F show kinetics curves for receptor interactions occurring intra-DOL, inter-DOL and in solution respectively. Overall, the interaction process is a trimolecular reaction where A^i^ or B^i^ intermediate is formed first as a bimolecular reversible process between a receptor and a reporter molecule (Supplementary Figure S2). Either intermediate can interact irreversibly with other complementary active receptor to form a ternary complex for which the rate constant was derived from a reaction between reporter complex and non-cholesterol modified receptors in solution (Figure 2F, note the receptor concentration is approximately two orders of magnitude higher than DOL cases in order to observe faster saturation kinetics; contrasting grey curve in Figure 4B with receptors at 5 nM). Using the model described in Supplementary Information Section 6 and Supplementary Figure S9, we deduce that, due to high local receptor concentration and constraints on a fluid surface, the effective rate constant of reaction is 2800-fold higher in DOL-bound receptors than that measured in the solution case. Our model fits very well considering 1A1B and 2A2B stoichiometries used in our DOL-based experiments.

**Figure 4:**
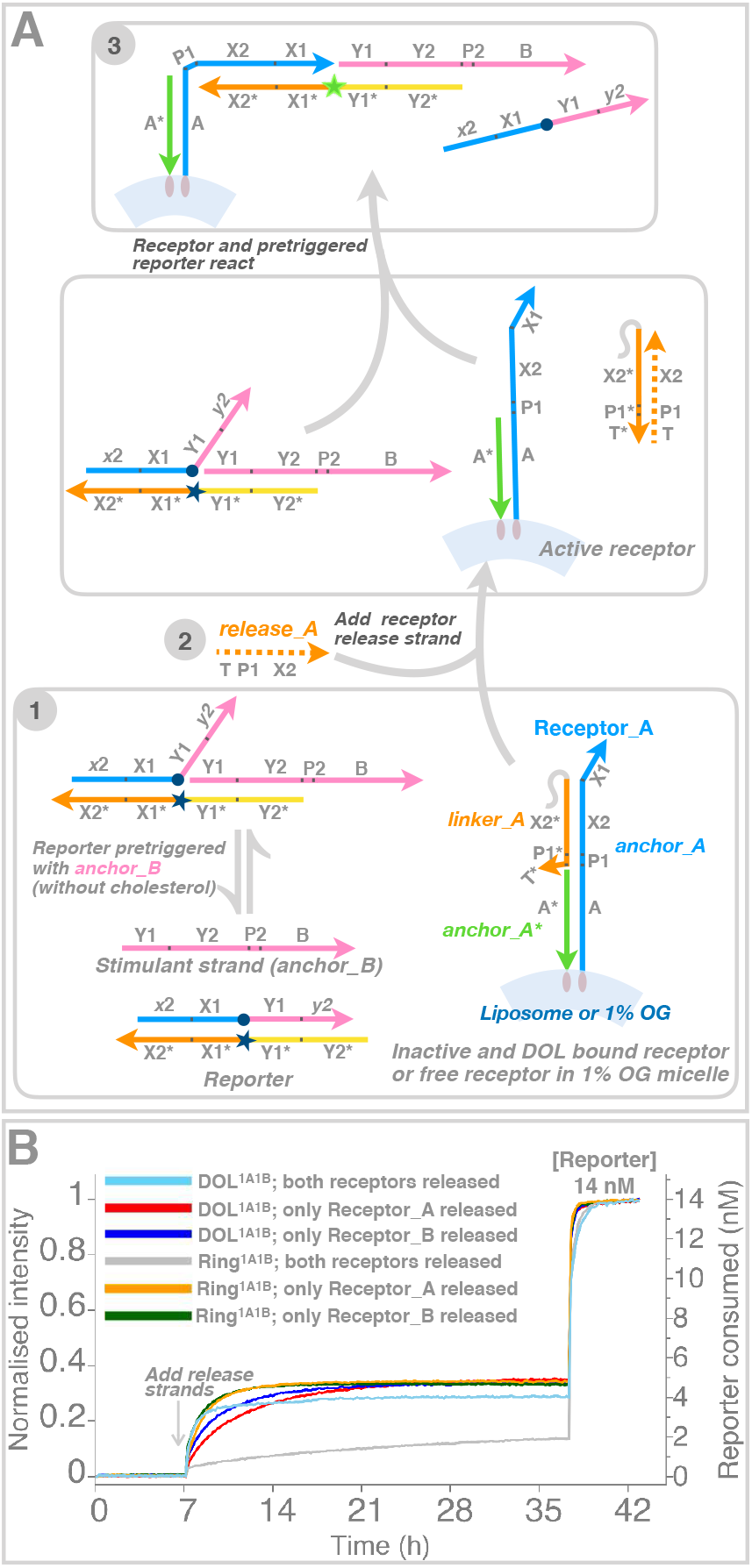
Determining tethering efficiency. Tethering efficiency of receptors to the DOL^1A1B^ platform was determined by comparing the extent of receptor reaction when both receptors were released and reacted normally within the DOL, and when one or the other receptor was reacted with a receptor complex in the presence of a stimulant strand in solution. **(A)** shows a modified logic circuit in which only receptor Receptor_A was released; a stimulant strand (*anchor_B* without a cholesterol modification) was supplied in excess to make up for any missing Receptor_B. A reciprocal experiment using *anchor_A* without a cholesterol modification is not shown. **(B)** Fluorescence curves (as in Figure 2) where either both the receptors were released with a normal reporter complex (cyan), only Receptor_A was released (red), or only Receptor_B was released (blue). Analogous curves are shown for a ring-only system (without a liposome), in which both the receptors were released (grey), only Receptor_A was released (orange, two repeats averaged), or only Receptor_B was released (green, two repeats averaged). As in Figure 2, reporter complexes were quenched after ~36 hours with an excess of both stimulant strands, or whichever was missing.

### Receptor tethering efficiency

Absolute stoichiometry control requires near 100% tethering efficiency of receptors. The DNA logic gate used for our DOL platforms is cooperative, requiring two different receptors to react with the reporter. If the tethering of receptors on the DOL ring is not 100% efficient, then it is possible to have four different DOL populations in the same purified fraction: DOL with no receptors, DOL with only Receptor_A, DOL with only Receptor_B, and DOL with both the receptors.

To evaluate tethering efficiency in DOL^1A1B^, we implement a DNA logic as shown in Figure 4A. The logic is similar to the logic shown in Figure 1B, but in this case only one receptor from DOL^1A1B^ platform was released while the other receptor remained tethered to the DNA scaffold. For example, the starting reaction mixture contained reporter complex (14 nM final concentration) with the purified DOL^1A1B^ along with an excess of stimulant strand (200 nM final concentration, a non-cholesterol version of *anchor_B*). The stimulant strand only partially triggers the reporter complex which does not completely displace the BHQ top strand. No rise in fluorescence was observed for the first 7 h (Figure 4B, red curve). After this, *release_A* (final concentration 200 nM) was added to selectively release Receptor_A which resulted in a sharp rise in fluorescence (red curve). A similar procedure was adopted to selectively release Receptor_B (blue curve), or both receptors at the same time (cyan curve).

Individually, completion levels of both the receptors, measured as a fraction of the total reporter complex consumed after all reporter is triggered, are about the same: ~ 4.9 nM for Receptor_A and ~ 4.7 nM for Receptor_B. Further, this indicates that the tethering efficiency is similar for both the receptors. While the concentration of DOLs with both receptors active is ~ 4.1 nM (completion level), assuming independence of tethering efficiency the total DOL concentration is about 5.8 nM (4.9×4.7 / 4). Thus, the calculated single labelling efficiency is 82-85%, and the double labelling efficiency is ~71%.

We also show an example where both receptors were tethered on a ring (5 nM, determined by absorption at 260 nm) without a liposome and were released together in solution (Figure 4B, grey curve) containing 1% n-octyl-β-D-glucoside (OG) detergent. The slower kinetics, in contrast to cases where at least one stimulant strand is in excess (yellow and green curves), is expected because here both the receptors are at only ~ 5 nM concentration. Interestingly, the saturation reached ~ 5 nM (almost same as ring concentration) in both the cases when either of the receptors was released. This could happen in a scenario where almost all the rings in the system have both the receptors, indicating high efficiency for the liposome-free system.

## Conclusions

Here we have shown stoichiometric, spatial and temporal (S^2^T) control for DNA receptor complexes in membranes for two different stoichiometries, which in principle could be scaled to different stoichiometries. A number of previous studies have examined the reaction of DNA receptors in membranes, either for the purpose of creating DNA circuits,^48–51^ studying diffusion within bilayers,^69^ or creating artificial signalling systems capable of transducing a DNA receptor dimerization event across a membrane.^70,71^ In particular, one study^49^ showed mild (75%) rate acceleration and significantly decreased leak for TMSD receptor reactions confined to liposomes, in the context of uncontrolled absolute stoichiometry. While none of these DNA receptor systems has achieved full S^2^T control, they provide inspiration for future uses of DOL. In the case of circuits, DOL will enable the implementation of systems where exact numbers of molecular inputs are required, or where each DNA computation cannot tolerate crosstalk with other copies of the DNA computation and must run within its own self-contained volume.^72^ And while we have demonstrated the release of up to four receptor reactants into the membrane, staple extensions on our current DOL could easily support the independently triggered release of several dozen different inputs, as required by a circuit, signalling cascade, or investigation of a biological question.

In our current approach, the receptors’ active domains (for release strand and reporter binding) are positioned between the linker to the ring and the hydrophobic groups (cholesterols) used as membrane anchors. This ensures that active domains are positioned outside of the liposome. On the other hand, signalling transduction systems^70,71^ suggest that it will be important to control the orientation of receptors inserted into DOL membranes, so that signal output domains can be positioned within the lumen of the DOL—intra-liposomally—when desired. In our system, an intra-liposomal domain could be added by (1) lengthening the hydrophobic groups so that they become a transmembrane domain and (2) attaching the desired domain to the distal end of the hydrophobic groups, so that it extends into the liposomal lumen. Ligands or auxiliary molecules meant to interact with intraliposomal domains could be either explicitly positioned with the same strategy, or simply encapsulated during the liposome formation. Where DOLs are used for membrane proteins, the position of the DNA linker (whether it is attached to the cytosolic or extracellular domain) will determine the orientation of the protein in the bilayer. When the linker is attached to the extracellular domain, the membrane protein is expected to be oriented “normally” so that the cytosolic domain is intraliposomal. When attached to the cytosolic domain the membrane protein will be “flipped”, with the cytosolic domain on the outside, where it can be studied and manipulated.

Each DOL is essentially a well-isolated reaction vessel with a controlled copy number of reactants—a digital nanoreactor. A bulk sample of DOL can therefore be measured without significant cross-reactions between vessels. As a result, properties which have until now required sophisticated single-molecule (or single liposome) techniques, can be measured using DOL via bulk fluorescence in a common plate reader. Detailed variation between reaction trajectories on different liposomes is, of course, averaged out by such bulk measurements, but variability in the number of molecules that can participate in a reaction is tightly controlled. In contrast, depending on the specific reactants and their concentrations, the extent of oligomerization and resulting size of aggregates can be unlimited in bulk experiments. As we have shown, bulk measurement of kinetics on DOL provides a sort of “integration over digital nanoreactors” that preserves kinetics as a function of copy number and maintains confinement of reactants to the restricted environment of the nanoreactor. Thus, while DOL could be examined with a single molecule technique, the DOL platform also enables a type of experiment whose window on the molecular world lies somewhere between that of a single molecule experiment and classical bulk technique (‘single molecule in bulk assay’).

We note that DNA nanostructure,^73,74^ DNA micelle,^75^ protein organelle,^76^ protein nanopore,^77^ viral,^78^ vesicle,^79^ MOFs^80^ and polymersome^81^ nanoreactors or zeptoreactors^82^ have been explored before, but none with the specific advantages provided by DOL. Viral capsids have encapsulated single enzymes^78^ and hollow DNA origami have encapsulated exact numbers of enzymes within a cascade^83,84^ but neither has yet enabled the exact number of reactants to be defined. DNA origami with reactants constrained to remain on their surface^34–38^ provide fully digital nanoreactors, with total control over the type and number of all reactants. Such membrane-free platforms have even stronger spatial control than do DOL, able to control local geometric configuration and reaction sequence. Especially interesting for applications in signal amplification, DNA computing^38^ and molecular robotics,^34–36^ they purchase extra spatial control at the cost of preventing reactants from diffusing freely within the nanoreactor, as occurs in our DOL platform.

With respect to diffusion of reactants within the DOL, several questions remain. Here we have not verified that the effective reaction area of the nanoreactors scales linearly with the membrane area of the liposome (e.g., by making larger or smaller liposomes). We have similarly not verified that receptors positioned away from the equator of the ring (say at opposite poles) exhibit similar behavior to those immobilized at the equator, to demonstrate the free diffusion of receptors from one hemisphere to the other (across the liposome’s zone of contact with the ring). Experiments to verify these aspects of DOL will be required to delineate the conditions under which DOL can be modeled as simple nanoreactors in which the membrane is homogeneous and its biophysical properties (e.g., receptor diffusion constant) are independent of DOL size. For proteins whose oligomerization behavior depends on membrane curvature,^85,86^ the assumption of DOL size-independent behavior will likely fail, making modeling more challenging. On the positive side, wherever membrane biophysics does turn out to be DOL size-dependent, development of a series of DOL having a range of diameters could enable new opportunities—e.g., protein sensors of membrane curvature could be studied and engineered.

Because a main motivation of this work is the eventual study and control of membrane protein interactions, it is important to discuss both the prospects and potential challenges. In principle, using DOL with membrane proteins should be as simple as replacing the DNA receptors with DNA-conjugated membrane proteins, where the protein-DNA linkers carry appropriate FRET probes. DOL are currently hybridized with cholesterol-modified DNA receptors in the presence of detergent, and so tethering detergent-solubilized membrane proteins (conjugated to appropriate DNA linkers) under similar conditions should be possible. However, as currently cast, the DOL system best models ligand or chemically induced protein interactions, where the reporter complex acts as the ligand to mediate receptor interactions. Such mediation by the reporter complex, as well as triggered activation of the receptor toeholds for the reporter complex by the release strands, provide two levels of protection against any receptor interaction before it is desired. The result is that the DOL are resistant against receptor-mediated inter-DOL leak reactions and DOL aggregation. In the case of ligand-induced protein interactions, where the proteins under study should have weak interactions before introduction of the appropriate ligand, we expect that current DOL will perform adequately.

In the case of proteins with constitutive interactions^87^ new techniques will be required to use DOL with minimal leak and aggregation; that is to keep proteins in their monomeric states. One approach may be to simply disrupt salt-sensitive constitutive interactions with high salt during synthesis and purification, before performing a concomitant trigger and buffer exchange step. Depending on the speed of intra-DOL versus inter-DOL reactions, this approach may be sufficient. For some proteins, whose interactions are denatured by detergent before the liposome forming step removes the detergent, orienting their oligomerization domains to the inside of the lumen may be sufficient. Overall, with just a few simple modifications in the basic technique, DOL digital nanoreactors may have the potential to provide custom instruments for the study and dissection of even the most complex membrane protein interactions.

## Materials and Methods

### Materials

Modified and unmodified DNA strands were purchased from Integrated DNA Technologies (IDT, USA). All staple strands, except those with linker extensions, were obtained and used in an unpurified form. Staples with *linker_A* or *linker_B* extensions were either purchased HPLC-purified or purchased unpurified and PAGE-purified in-house before use. All receptor and reporter complex strands were purchased HPLC-purified, dissolved in 1×TE buffer and stored at −20 °C. Sequences for cholesterol-modified DNA (with a triethylene glycol linker), including IDT modification codes are provided in Supplementary Table S1. Lipids were purchased from Avanti Polar Lipids, USA. Gels were imaged using a ChemiDoc MP instrument (Biorad, USA). In many buffers n-octyl-β-D-glucoside (OG) was added as a detergent. Origami annealing buffer is 1xTE, 12.5 mM MgCl_2_; TE-Mg buffer is 1xTE, 10 mM MgCl_2_; TAE-Mg buffer is 1×TAE, 10 mM MgCl_2_; TE-Mg-OG buffer is 1% OG, 1×TE, 10 mM MgCl_2_; HEPES-Mg-K is 10 mM MgCl_2_, 25 mM HEPES, 100 mM KCl; HEPES-OG buffer is 1% OG in HEPES-Mg-K; in all cases 25 mM HEPES buffer pH 7.4 adjusted with KOH. Where possible, final concentrations (f.c.) of solution components are given.

### Ring design, assembly and purification

We used a DNA origami ring design reported earlier^59^ with slight modifications for positioning linker strands. caDNAno^88^ designs and staple sequences are provided in the Supporting Information. DNA scaffold (8064 nucleotides), 100 nM f.c., was mixed with 6x excess of staple strands, including linker strands in origami annealing buffer. DNA scaffold was produced from *E*. *coli* and M13 derived bacteriophages.^29^ Typically, 1000 μL reaction mix (scaffold and staples) was prepared and divided in 20 tubes. All tubes were annealed from 95 to 20 °C over 36 h and then the annealed reactions were pooled and concentrated using 30 kDa Amicon 0.5 mL centrifugal filters. Filters were pre-wetted with TE-Mg by centrifuging at 6000 RCF for 4 minutes. Afterward, pooled annealed reaction mix was concentrated by loading 500 μL volume in two different filters by centrifuging at 8000 rpm for 8 minutes. Concentrated sample (total ~180 μL) was mixed with glycerol (f.c. ~7%) and divided in two equal volumes for further purification. To make a gradient, ~2.5 mL each of 15% and 45% glycerol in TE-Mg were loaded initially into an ultracentrifuge tube to form two layers, which were converted into a continuous gradient using Biocomp gradient station. Finally, each volume (in 7% glycerol mentioned above) was loaded on top of freshly made gradient and purified using rate-zonal ultracentrifugation by rotating at 304,000 RCF for 1 h at 4° C. After this, ~20 fractions (200 μL each) were collected manually from the centrifuge tubes. To determine the fraction containing desired product, 5 μL of each fraction was loaded in 1.5 % agarose gel (prepared with TAE-Mg buffer having ethidium bromide as a pre-stain) and the gel was run at room temperature by applying 60 V for 1.5 h in TAE-Mg. Based on gel results (Supplementary Figure S1B) the desired fractions were pooled and concentrated using 30 kDa Amicon 0.5 mL centrifugal filters (as above). At the end of this step, only trace amounts of staples remained. To remove glycerol from the concentrated sample, we performed one or two 400 μL TE-Mg washes; trace glycerol at this step did not affect downstream steps. Ring concentration was by measuring UV absorption at 260 nm using a Nanodrop spectrophotometer. Purified rings were stored at 4 °C (and used within a week) or −20 °C (and used within one or two months).

### Annealing reporter and receptors

For the reporter complex, top strand (with black hole quencher, see Table 1 for IDT order code) was added in 1.5× excess of the bottom strand (with fluorescein, see Table 1 for IDT code) with f.c. 300 nM and 200 nM respectively. The total volume in TE-Mg buffer was ~1000 μL. Reaction mix was annealed in different tubes (each ~100 μL) from 95 to 20 °C over 2 h. Annealed reactions were pooled together, stored at 4 °C, and later used without further purification. The same batch of reporter complex was used for all plate reader measurements. Freshly thawed and annealed volumes of cholesterol receptor complexes were used for each experiment. 10 μM aliquots of the cholesterol-modified strands stored at −20 °C were thawed at room temperature at least for 1 h. Annealing was performed from 95 to 20 °C over 2 h using 2x excess of *anchor_A*\* or *anchor_B*\* (f.c. 600 nM) with *anchor_A or anchor_B* (f.c. 300 nM) in TE-Mg-OG. Annealed receptors were used further without purification.

### Tethering DNA receptors to rings and purification

Purified rings containing linkers were incubated with freshly annealed receptors at 37° C for 1 h in TE-Mg-OG buffer modified to have 1.15% OG. For DOL^1A1B^, Receptor_A and Receptor_B (f.c. 90 nM each) were added at 3× in excess of ring (f.c. 30 nM) containing one *linker_A* and one *linker_B*. For DOL^2A1B^, Receptor_A (f.c. 135 nM) was 4.5× in excess while Receptor_B was 3× in excess of ring (f.c. 30 nM) containing two *linker_A* and one *linker_B*. For DOL^2A2B^, Receptor_A and Receptor_B (f.c. 135 nM each) were 4.5× in excess of ring (f.c. 30 nM) containing two *linker_A* and two *linker_B*. In general, the total incubation volume was ~200 μL. To remove the excess receptors and to determine the desired fractions, we followed a rate-zonal ultracentrifugation purification procedure and agarose gel analysis steps similar to those described above for rings, with minor differences. Here, a 15–45% glycerol gradient was prepared with detergent (in TE-Mg-OG) and centrifuged at 10° C (rather than 4° C). Desired fractions were pooled and concentrated using 30 kDa 0.5 mL Amicon centrifugal filters, with one or two 400 μL final TE-Mg-OG buffer washes. Ring concentration was estimated by UV absorption at 260 nm using a Nanodrop spectrophotometer; the purified product was stored at 4° C and used the next day.

### DOL formation and purification

Stock 10 mM lipid mixture was made with 75:20:5 molar ratio of 1,2-dioleoyl-sn-glycero-3-phosphocholine (DOPC), 1,2-dioleoyl-sn-glycero-3-phospho-L-serine (DOPS), 1,2-dioleoyl-sn-glycero-3-phosphoethanolamine-N-[methoxy(polyethylene glycol)-2000] (PEG2000-PE) respectively in chloroform (f.c.: 7.5 mM DOPC, 2 mM DOPS, 0.5 mM PEG2000-PE). A desired volume of this stock was dried under nitrogen gas for 10**–**20 min and then further dried for 3 h in a freeze dryer (Freezone 1, Labconco). For use, dried lipids were rehydrated to a concentration of 10 mM lipids with 25 mM HEPES and 100 mM KCl buffer and shaken for 0.5 h at room temperature. The ring has handles (32 staple extensions in the case of two receptors and 30 for four receptors), which can hybridize with antihandles made of cholesterol-modified oligonucleotides (Step C, Figure 1A). These antihandles act as seeds for liposome formation. Each purified sample of ‘rings with hybridized receptors’ (f.c. 30 nM) was incubated with cholesterol-containing antihandles (f.c. 1.8 μM) at 37 °C for 1 h in HEPES-OG buffer. After incubation, each sample of ‘rings with hybridized receptors and antihandles’ (f.c. 15 nM) was mixed with hydrated lipids (f.c. 1.5 mM) in HEPES-OG buffer to create a total volume ~150 μL and was shaken gently for 0.5 h at 25 °C. To remove the detergent and to form liposomes inside the rings, the mixture was transferred to Slide-A-Lyzer 0.5 mL 7 kDa dialysis cassette using a syringe. Dialysis was done overnight at room temperature against 2 L HEPES-OG buffer.

To purify the dialysis mix we performed isopycnic ultracentrifugation, using 6–30% iodixanol gradients in HEPES-Mg-K where less dense free liposomes float to the top, and rings holding liposomes are distributed in lower fractions. After overnight dialysis we typically recovered ~210 μL per sample. For each sample, 200 μL was used and divided in two 100 μL replicates and each replicate was mixed with 200 μL of 45% iodixanol in HEPES-Mg-K. Thus for each replicate a total of 300 μL containing 30% iodixanol was placed at the bottom of an ultracentrifuge tube, above which 60 μL each of 26%, 22%, 18%, 14%, 10%, and 6% of iodixanol were layered (bottom to top) via manual pipetting. Samples were centrifuged at 280,000 RCF for 5 h at 4 °C and twelve or thirteen 50 μL fractions were collected from each centrifuge tube. Fractions were collected in tubes that had been pre-rinsed with a blocking solution (1 μM 15T oligonucleotides in HEPES-Mg-K buffer); all tubes used after this step (for pooling or transfer) are also pre-rinsed with blocking solution. For each DOL, identical fractions from replicates were pooled, and pooled fraction 3 and fraction 4 were further combined. To each pooled sample 15T oligo was added to 1 μM f.c.

### Fluorescence plate reader experiments

A Biotek Cytation-1 plate reader was used for real-time fluorescence measurements. Plate reader measurements were done at 25 °C using a 475/20 nm excitation filter and a 530/25 nm emission filter. Samples were loaded manually into Corning 384-well assay plates (black with clear flat bottoms). To avoid sample evaporation, plate wells were sealed with Nunc polyolefin acrylate sealing tape. Before loading samples, wells were pipette-rinsed with blocking solution. To each DOL tested, reporter complex was added (4.7 or 14 nM f.c.) and samples were mixed gently via manual pipetting. Next, 46.2 μL of each sample was loaded per well, making sure no air bubbles were trapped in the wells. Baseline fluorescence was first measured for ~7 h. Release strands were added (0.9 μL of a stock containing 5 μM each of *release_A* and *release_B* to create 100 nM f.c. of each release strand) to initiate receptor interactions, which were measured for a further ~18 h. To establish a maximum fluorescence endpoint, with which each samples trace could be normalized, we triggered any remaining reporter complex by adding excess *anchor_A* and *anchor_B* strands (versions without cholesterol modifications, to 100 nM f.c. for each) and then measured the fluorescence for another 4 to 6 h.

### TEM sample preparation

Uranyl formate negative stain solution (1% w/v) is acidic and can denature DNA nanostructures; thus one ml aliquots were neutralized by adding 2.5 μl of 5M NaOH prior to use (see guidelines for preparation and storage elsewhere^89^). DOL samples (5 μL) were deposited on a glow-discharged formvar/carbon coated copper grid (Ted Pella, Inc.) for 1 minute, and liquid was blotted away using filter paper. Each grid was subsequently washed with 7.5 μL of HEPES-Mg-K buffer and stained with 7.5 μL neutralized uranyl formate negative stain for 1 minute. Negative-stain TEM images were acquired using an FEI Tecnai T12 TEM (120 kV) equipped with an EDS detector and 4k x 4k Gatan Ultrascan CCD.

### Analyses and plots

Raw plate reader data (in text format) analyses were performed using custom PERL scripts. Plots were created using XMGRACE^90^ or *gnuplot* (www.gnuplot.info).

## Supporting information

Supporting info for main manuscript

caDNAno design and staple sequences

## Author contributions

V.M. conceived the original idea, designed, performed and analyzed most experiments, and wrote the manuscript first draft. V.M., Z.Z., C.T. and P.W.K.R. contributed further ideas. C.T., V.M., and P.W.K.R. designed the DNA logic circuit and analyzed kinetics data. Z.Z. designed the origami, assisted in DOL synthesis, and performed TEM. N.S. and C.T. modelled the DNA circuit. E.R.C. mentored and hosted V.M. All authors discussed the results and participated in manuscript writing.

## Acknowledgements

V.M. acknowledges Erik Winfree and Lulu Qian for both their comments and access to equipment for initial experiments. V.M. thanks Human Frontier Science Program (HFSP LT001164/2017-L) for a Postdoctoral Fellowship and Caltech for additional support. This study was supported by National Institute of Mental Health awards MH125320 (to P.W.K.R. and E.R.C.), MH061876 and NS097362 (to E.R.C.), and a Faculty Early Career Development Award from NSF CCF 2143227 (to C.T.). E.R.C. is an investigator of the Howard Hughes Medical Institute (HHMI). This article is subject to HHMI’s Open Access to Publications policy. HHMI lab heads have previously granted a nonexclusive CC BY 4.0 license to the public and a sublicensable license to HHMI in their research articles. Pursuant to those licenses, the author-accepted manuscript of this article can be made freely available under a CC BY 4.0 license immediately upon publication.

## Competing interests

The authors declare no competing financial interests.

## Notes

### Competing Interest Statement

The authors have declared no competing interest.

