## Supporting info for main manuscript for "Digital nanoreactors for control over absolute stoichiometry and spatiotemporal behavior of receptors within lipid bilayers"

### Supplementary information

#### 1) Analysing different fractions

We performed initial experiments to determine which fractions contain DOL among the different fractions collected after isopycnic purification step (Step D Figure 1A and Supplementary Figure S1D). We confirm this by transmission electron microscopy (TEM) images and fluorescence measurements using a plate reader instrument on different fractions for DOL$^{1A1B}$ case (Figure3 and Supplementary Figure S3). TEM data suggested that fractions 1 and 2 have only free liposomes, and the count of free liposomes is higher in fraction 1 than fraction 2. Most of the DOL were found to be present in fractions 3-5 and rarely observed beyond fraction 6. TEM images for DOL clearly show a DNA ring structure around a spherical blob.

To corroborate TEM observations we performed plate reader experiments. Ideally, assembly and purification steps explained in Figure 1A should result in a DOL platform with the tethered receptors (mainly found in fractions 3-5 as observed in TEM) without any free diffusing receptors either in the solution or on the DOL surface. It is more likely that free cholesterol receptors, if there are any, should anchor in the membrane surface of free liposomes (fractions 1 and 2) or DOL lipid bilayer (fractions 3-5) rather than diffusing in the bulk solution. It is important to note that free receptors are active and both the receptors can cooperatively displace the top quencher strand from the reporter complex which could result in undesired fluorescence signal. Keeping these factors in mind we tested the logic circuit in all the collected fractions (up to 10). As shown in Figure 3 and Supplementary Figure S3, we started by mixing each fraction with the reporter complex (final concentration 4.7 nM) and for initial 7 h we did not observe any major fluorescence signal in any fraction. If there were any free receptors, in any fraction, then it would have resulted in a rise in signal over the initial 7 h. After 7 h, both release strands were added which released both the receptors (thus active) and their cooperative hybridisation with reporter complex resulted in a sudden rise in fluorescence signal. The saturation in a signal (amount of reporter consumed) is proportional to the receptor concentration; thus the DOL concentration which contains the receptors. Both the TEM data as well as fluorescence data correlate very well. TEM data suggested that fractions 3 and 5 have lower DOL concentration (based on visual counts) than fraction 4. Similarly, the fluorescence signal obtained from fraction 3 and 5 is lower than the fraction 4. Based on this data, for our other experiments we decided to combine fractions 3 and 4 to have more volume in hand to perform repeats and other controls. In contrast, the highest concentration of DOL in *dimer*_DOL$^{1A1B}$ case was found to be in fraction 5 (Supplementary Figure S7) instead of fraction 4 as in the case of DOL$^{1A1B}$, which is expected because *dimer*_DOL$^{1A1B}$ has two rings dimerised together (Supplementary Figure S6).

#### 2) Miscellaneous note

Our purification step (Step A, Figure 1A) removes most of the staples, including linkers, and further the purified fractions (~ 97% origami band intensity, ~ 3% traces staples band intensity, Supplementary Figure S1B) were pooled and concentrated using Amicon 30 kDa centrifuge filters (Materials and Methods) which should additionally get rid of traces of staples or linkers. Note that ~3% band intensity is for all the staples (~200 in number) and the contribution from linker strands (2-4 in number) would be almost negligible (if contribution is equally weighed among all strands then it would be < 0.02%). If we assume, which is less likely, the traces of linker strands still remain around after Step A purification and the filter concentration step (Figure 1A) then the free linkers may hybridise with the receptors in solution forming traces of 'protected' receptors at Step B (Figure 1A). If we further assume that traces of protected receptors were not removed at all, which is again unlikely, in further next steps (Step B purification) then at least one can say all of the signal (curve kinetics) that we see for DOL after the release strand is added is originating from > 99% of tethered receptors and rest is the undesired reaction coming from protected receptors on the DOL surface.

**3) Concentration from saturation end points**

For the cases (main Figures 2D and 4B, Supplementary Figures S4A, S5, and S7) where the reporter complex is not fully consumed, the saturation end points could be used to determine the DOL concentration. This requires measurement of maximum available fluorescence to know the amount of unreacted reporter, which we performed in all the cases by adding excess of stimulant strands (non-cholesterol versions of *anchor_A* and *anchor_B*). Before explaining how we used this method to determine DOL concentration, it is important to note that adding excess of a reporter complex does not occupy/block free individual receptors (partial displacement of BHQ top strand by free single receptor; $A^i$ and $B^i$ states shown in Supplementary Figure S2). To justify this, as shown in Supplementary Figure S5, when the same $DOL^{1A1B}$ sample is mixed with two different reporter concentrations, 4.7 nM and 14 nM (*c.a.* 3 times higher) in both the cases the kinetic curves end almost at the same saturation point after the receptors are released to react with the reporter; implying that same amount of reporter complex is consumed. If excess reporter were to occupy both the receptor types individually, in the form of favorable stable intermediate states ($A^i$ and $B^i$), then the end points (reporter consumed by receptors) reached using two different reporter concentrations would not have been the same leaving some receptors incapable to react cooperatively (saturation level in case of 14 nM reporter concentration expected to be lower in such a scenario). Additionally, the final end product, a ternary complex showing fluorescence, is thermodynamically more favourable than the intermediate states ($A^i$ and $B^i$), which makes the forward process almost irreversible. Overall, this implies that the receptor and reporter molecules would be consumed irreversibly in the form of a final ternary complex only (without any stable intermediate states) and yielding a fluorescence signal. In our experiments the release strands are added in excess (100 nM), which makes TMSD receptor release almost complete and ensures that the receptors on every DOL are available and active to react with reporter complex in solution. To summarize this, for the cases where receptor concentration is lower than reporter concentration we used saturation end points to determine the DOL concentration.

Furthermore, we can counter the argument that we do not know whether release strands were in excess or not because we do not know DOL concentration in any fraction (each 50 μL), by pointing out that the release strands (100 nM) are indeed in excess of the starting raw material itself before purification (ring at 15 nM, 150 µL at Step D, Figure 1A). Further supporting this point is the fact that we used a known concentration, 5 nM of a purified ring tethered with one Receptor_A and one Receptor_B (obtained after Step B, Figure 1A), and then quantified the concentration of each receptor independently by releasing one receptor at a time (Figure 4B; orange and green curves). This data showed that the two receptors were at almost equal concentrations and their individual concentrations were almost equal to the ring as well (5 nM).

Therefore, considering the discussion above, we can assume that the saturation end point is simply not the measure of concentration of species in equilibrium but is an indication that the reaction goes to completion consuming almost all the available receptors on each DOL. This is the rationale we employed in using saturation end points as the measure of DOL concentration as shown in Figure 2D; [$DOL^{1A1B}$] is ~2.4 nM, [$DOL^{2A2B}$] is ~1.4 nM and [*dimer_* $DOL^{1A1B}$] is ~1.2 nM. Note that in the case of $DOL^{2A2B}$, the DOL platform concentration (for fraction 5, Figure 2D) would be half of the receptor concentration (saturation point), where receptor concentration is equal to the reporter consumed. Concentration of $DOL^{2A2B}$ in a combined fraction (3+4) cannot be evaluated from the curve saturation end points as all the reporter molecules were consumed (Supplementary Figure S4A); at 36 h adding excess of stimulant strands did not show any further spike in the signal but receptor concentration is at least 4.7 nM. Nonetheless, we indirectly estimated $DOL^{2A2B}$ concentration in the combined (3+4) fraction using TEM data (Supplementary Section 4) and found it to be *c.a.* 2.98 ± 0.53 nM, thus making receptor concentration *c.a.* 6 nM. Thus, these results are in good agreement with the plate reader observation that the combined (3+4) fraction consumed all the reporter molecules (at 4.7 nM).

**4) Concentration from TEM**

As discussed in Section 3 above the kinetic curves, Figure 2D, show that the reporter complex is not fully consumed in these cases and thus saturation end points could be used as a measure of DOL concentration. We also collected TEM images for $DOL^{1A1B}$ samples which allowed us to correlate the number of counts of DOL (done manually) with the concentration obtained from the plate reader experiment. Linear regression fitting of concentration (from plate reader) of $DOL^{1A1B}$ *vs*. their averaged TEM counts (taken from at least 4 TEM images for $DOL^{1A1B}$ case from two different fractions) provided a general relationship between counts and concentration (Supplementary Figure S4B bottom). This allowed us to calculate concentration of pooled fraction (3+4) in case of $DOL^{2A2B}$ to be ~2.98 nM (green curve Supplementary Figure S4A, where all the reporter complex was consumed) from TEM data (averaged manual counts from 7 images of pooled fraction '3+4' $DOL^{2A2B}$ sample). Similarly, for inter-DOL cases (concentrations and plots in Figure 2E), which did not reach saturation, their concentrations were calculated using the same method described here (averaged manual counts from 5-8 TEM images in interDOL cases). Calculated concentrations for interDOL cases are: ~ 1.26 nM $interDOL^{1A1B}$ case and ~ 1.03 nM $interDOL^{2A2B}$ case.

**5) Dimer DOL**

Here a slightly different type of platform is discussed. As a recall, the main motive to develop DOL platform is to start with a configuration where monomeric forms of the molecular species in question (synthetic receptors or proteins) are well separated before assaying their interaction. The DOL platform discussed in the main paper (Figure 2A,B) would be suitable to assay chemically induced protein interaction, for example the homo- or hetero- dimerization of proteins in the presence of a ligand (drug, ion, peptide, soluble protein *etc.*) found in many protein pairs.[4] Considering our protocol step where we incubate, in a single pot, DNA origami ring with both the receptors (Step B, Figure 1A) it would not be suitable for proteins which could possibly form oligomers without requiring any ligand (proteins with constitutive interactions). For a chemical induced dimerization process this would not be an issue because at a specific tethering site there will be only one type of monomer present. For constitutive proteins this protocol could be problematic because there could be already an oligomeric form of protein complex attached at a specific tethering site. Though one could potentially test different conditions (*e.g.* detergents, salt concentrations, inhibitors *etc*. which can be later dialysed out at Step D, Figure 1A) to prevent oligomerisation at the incubation step but to overcome this we devised a slightly different strategy. We incubated separately each type of receptor with a DNA ring having a specific complementary linker and purified them individually (Supplementary Figure S6). Later, two DNA rings (each with only one type of receptor; Receptor_A or Receptor_B) were dimerised with the help of 16 complementary sticky staple end extensions on one ring to the scaffold of second ring. To assemble a liposome inside the dimerised rings (*dimer*_DOL$^{1A1B}$) we further followed the same protocol employed for DOL platforms. Similarly, receptors were released in *dimer*_DOL$^{1A1B}$ case and the kinetic curve for fraction 6 is shown Figure 2D (green curve). Supplementary Figure S7 shows the kinetic curves for different fractions using reporter complex at two different concentrations. Though for our *dimer*_DOL$^{1A1B}$ we still use chemically induced dimerization (where DNA receptors dimerise in the presence of reporter) we hope that similar strategy could be used for constitutive proteins as well.

**6) Kinetics simulation**

The reactions in this work were simulated using mass-action kinetics. Each reaction was modelled as a differential equation which was then solved by using a CRN Simulator Package.[1]

There are several reaction steps, listed below.

1) $release\_A + Receptor\_A(bound) \xrightarrow{K_{eff}} Receptor\_A(free) + release\_A{:}\,linker\_A$

2) $release\_B + Receptor\_B(bound) \xrightarrow{K_{eff}} Receptor\_B(free) + release\_B{:}\,linker\_B$

3) $Receptor\_A(free) + Reporter \underset{K_b}{\overset{K_f}{\rightleftarrows}} Receptor\_A{:}\,Reporter$

4) $Receptor\_B(free) + Reporter \underset{K_b}{\overset{K_f}{\rightleftarrows}} Receptor\_B{:}\,Reporter$

5) $Receptor\_A{:}\,Reporter\ +\ Receptor\_B(free) \xrightarrow{x.K_{eff2}} TernaryComplex\ +Top\ strand$

6) $Receptor\_B{:}\,Reporter\ +Receptor\_A(free) \xrightarrow{x.K_{eff2}} TernaryComplex\ +\ Top\ strand$

In reactions (1) and (2), *release_A* and *release_B* strands bind to their respective receptors in order to free them from the linker strands. The strand displacement reaction is irreversible and described by rate $\boldsymbol{K_{eff}}$, which is the effective rate of strand displacement. In reactions (3) and (4), the free receptors bind to the reporter. The forward rate, $\boldsymbol{K_f}$, is the rate of hybridization of the toehold; the backward rate, $\boldsymbol{K_b}$, is the rate of dissociation, which is determined by the formula[2] $10^{6-L}\ M^{-1}s^{-1}$, where L is the toehold length, in this case L = 5. In reactions (5) and (6), the receptor:reporter complex binds to the other receptor and does strand displacement at the rate $\boldsymbol{K_{eff2}}$, which describes the rate of cooperative strand displacement. This rate was determined using simulations of the reaction system in solution (using strands without cholesterol modifications, Figure 2F). The factor *x* is used to describe the speed-up in the reaction rate between the solution case and the DOL case, which in turn gives us insight into the relative concentrations of the receptors when they are on the DOL. This becomes clear when we look at the differential equations for that set of reactions:

(7) $$\frac{d[TernaryComplex]}{dt} = x * K_{eff2}\ [Receptor\_A{:}\,Reporter]\ [Receptor\_B(free)]$$

The rates used are[2,3]:

$\boldsymbol{K_{eff}} = 2*10^{5}\ M^{-1}s^{-1}$ [ref[2]]

$\boldsymbol{K_f} = 2*10^{6}\ M^{-1}s^{-1}$ [ref[3]]

$\boldsymbol{K_b} = 10\ s^{-1}$

$\boldsymbol{K_{eff2}} = 7.6*10^{4}\ M^{-1}s^{-1}$

Experimental curve fittings based on this model are shown in Supplementary Figure S9.

#### 7) DNA ring origami design and staple sequences

DNA origami ring design (caDNAno json file) and staple sequences (including linkers for receptors) are provided as supplementary information files. Nanocage_BASIC.json is a general design and the staple sequences for a specific design can be found with their helix and residue numbers (caDNAno format) in *.csv* files. For dimerization of rings in dimer_DOL[1A1B] case there are 16 staples extended (labelled as 'dimer-linker' in file *Ring2_for_dimer1A1B_sequence.csv*) with small sticky ends (4 nucleotides) from one ring which hybridize with the scaffold of the second ring (some staples from second ring are thus kept a bit shorter).

In all cases handle sequence for cholesterol-antihandle (extension from staples):
`AAATTATCTACCACAACTCAC`
In all cases antihandle sequence with cholesterol (HPLC purified):
`/5Chol-TEG/GTGAGTTGTGGTAGATAATTT`

**Table S1**: Domain decomposition of DNA receptors (see Figure 1). Domain names, their sequences (5'—3') and their roles are summarised. Colour codes used here correspond to the same coloured domains shown in Figure 1.

| Domain | Sequence (nucleotide length) | Role |
|---|---|---|
| **Receptor_A complex** (*anchor_A*, *anchor_A**) | | |
| **A*** (*anchor_A**) | GTTTGAGTTGAGTGGGAAAG/3CholTEG/ (20) | Cholesterol anchor |
| **A** (*anchor_A*) | /5Chol-TEG/CTTTCCCACTCAACTCAAAC (20) | Cholesterol anchor |
| **P1** (*anchor_A*) | CA (2) | *linker_A* tethering, unpaired hinge in ternary complex |
| **X2** (*anchor_A*) | ACACCATTTACCCAC (15) | *linker_A* tethering, reporter toehold binding |
| **X1** (*anchor_A*) | ATTCAAATCC (10) | Cooperative hybridisation (along with *X2*, *Y1Y2*) |
| ***linker_A*** (extension of staple strand connected *via* TTTT spacer) | | |
| **X2*** | GTGGGTAAATGGTGT (15) | Receptor_A complex tethering |
| **P1*** | TG (2) | Receptor_A complex tethering |
| **T*** | AGATG (5) | Toehold for *release_A* |
| **Receptor_B complex** (*anchor_B*, *anchor_B**) | | |
| **B*** (*anchor_B**) | /5Chol-TEG/GTTGGTAATGGAATGGGAAG (20) | Cholesterol anchor |
| **Y1** (*anchor_B*) | CACAATACAC (10) | Cooperative hybridisation (along with *Y2*, *X1X2*) |
| **Y2** (*anchor_B*) | CCTACACATACATCA (15) | *linker_B* tethering, reporter toehold binding |
| **P2** (*anchor_B*) | AC (2) | *linker_B* tethering, unpaired hinge in ternary complex |
| **B** (*anchor_B*) | CTTCCCATTCCATTACCAAC/3CholTEG/ (20) | Cholesterol anchor |
| ***linker_B*** (extension of staple strand connected *via* TTTT spacer) | | |
| **S*** | GTGGA (5) | Toehold for *release_B* |
| **P2*** | GT (2) | Receptor_B complex tethering |
| **Y2*** | TGATGTATGTGTAGG (15) | Receptor_B complex tethering |
| ***Releasing strands*** | | |
| **T** (*release_A*) | CATCT (5) | *linker_A* toehold binding and strand displacement |
| **P1** (*release_A*) | CA (2) | |
| **X2** (*release_A*) | ACACCATTTACCCAC (15) | |
| **Y2** (*release_B*) | CCTACACATACATCA (15) | *linker_B* toehold binding and strand displacement |
| **P2** (*release_B*) | AC (2) | |
| **S** (*release_B*) | TCCAC (5) | |
| ***Reporter complex*** | | |
| **x2.X1.Y1.y2** *(Top strand with quencher)* | atttacccac.ATTCAAATCC./**iBHQ-1dT**/.CACAATACAC.cctacacata (10.10.1.10.10) **iBHQ-1dT** is black hole quencher, internal modification on T | |
| **Y2*. Y1*. X1*. X2*** *(Bottom strand with fluorophore)* | TGATGTATGTGTAGG.GTGTATTGTG. /**iFluorT**/.GGATTTGAAT.GTGGGTAAATGGTGT (15.10.1.10.15) **iFluorT** is fluorescein, internal modification on T | |

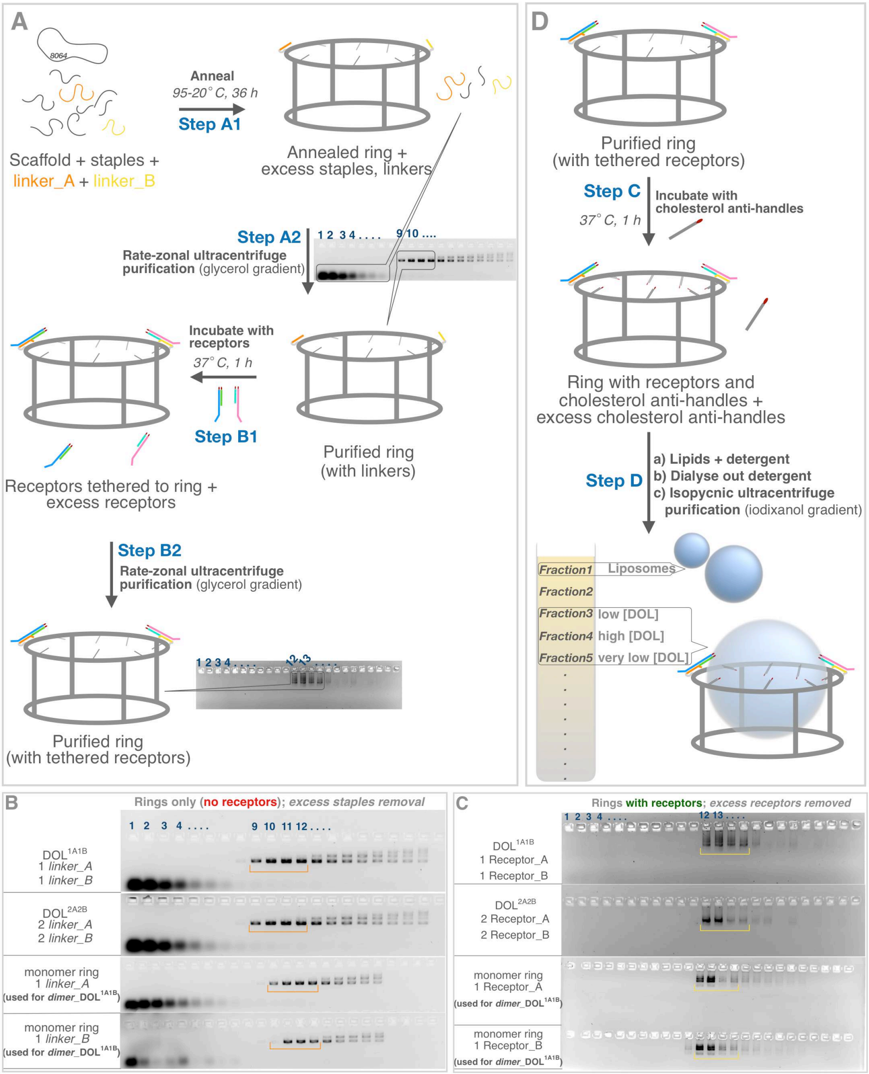


**Figure S1**: *DOL platform assembly and purification*. Steps in Figure 1A are elaborated here for different DOL cases. **(A)** After annealing (Step A1) excess staple strands were removed (Step A2) by rate-zonal ultracentrifuge purification. Different fractions so collected were run through 1.5% agarose gel shown in **B**. Fractions containing (9-12 shown in **B**) the purified rings were pooled and concentrated. After after incubation with receptors (Step B1) excess receptors were removed by rate-zonal ultracentrifuge purification and corresponding agarose gels are shown in **C**. Fractions containing (12-14 shown in **C**) the purified rings with tethered receptors were pooled and concentrated. **(D)** Elaborated Steps C and D in Figure 1 are shown here. Last step is shown to give an idea about the distribution of liposome and DOL in the tube after isopycnic separation.

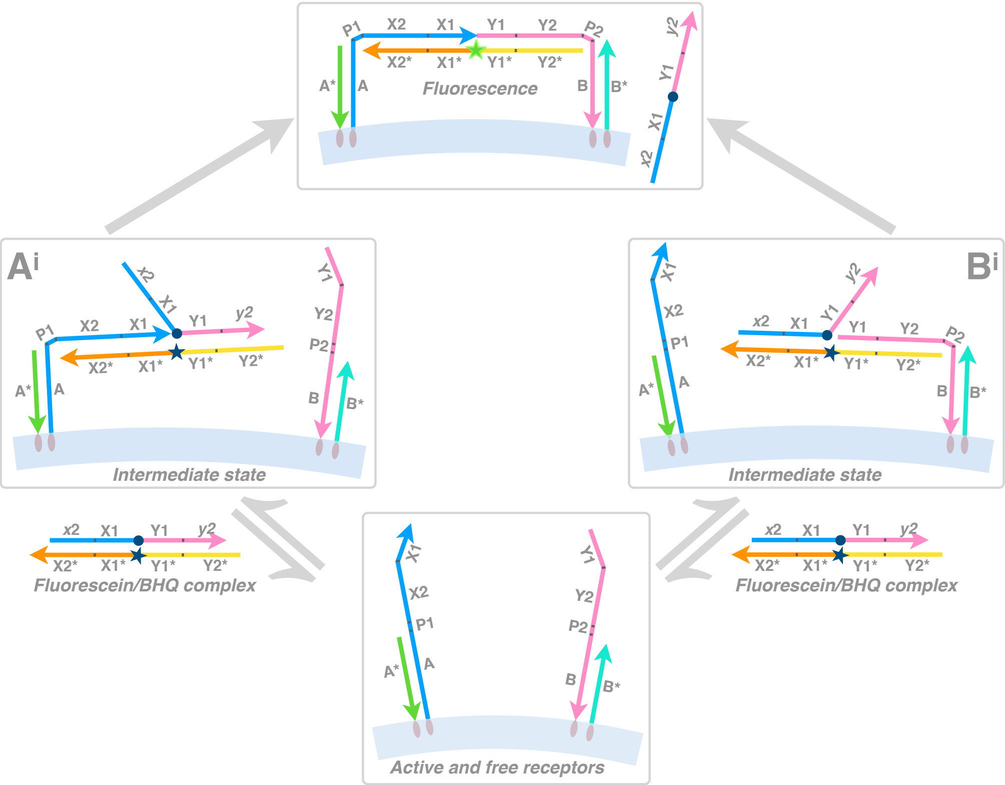


**Figure S2**: *Intermediate states leading to ternary complex*. Possible intermediate states are shown here for the DNA circuit in Figure 1B. Lower case domains are partially complementary to their upper-case counterparts, *e.g.* **x2** is a shortened version of **X2** and is only partially complementary to **X2***.

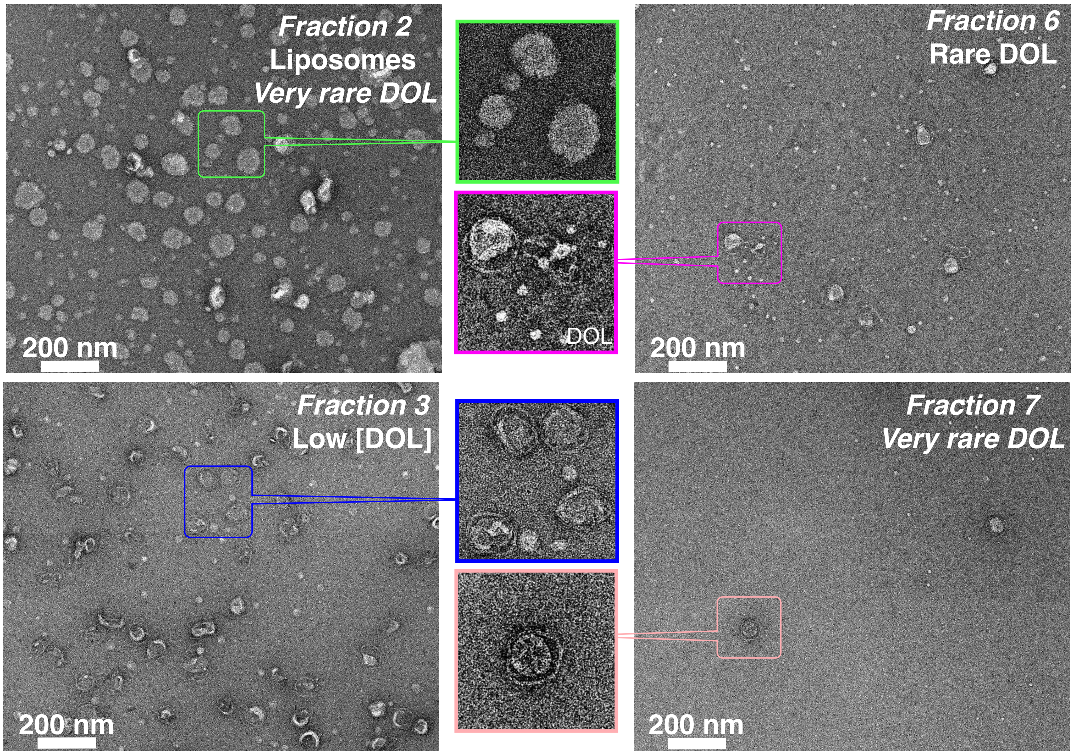


**Figure S3**: *Analyzing different fractions*. TEM images shown for fractions 2, 3, 6 and 7 related to Figure 3 (which shows fractions 1, 4, and 5). None of these fractions have any significant amount of DOL.

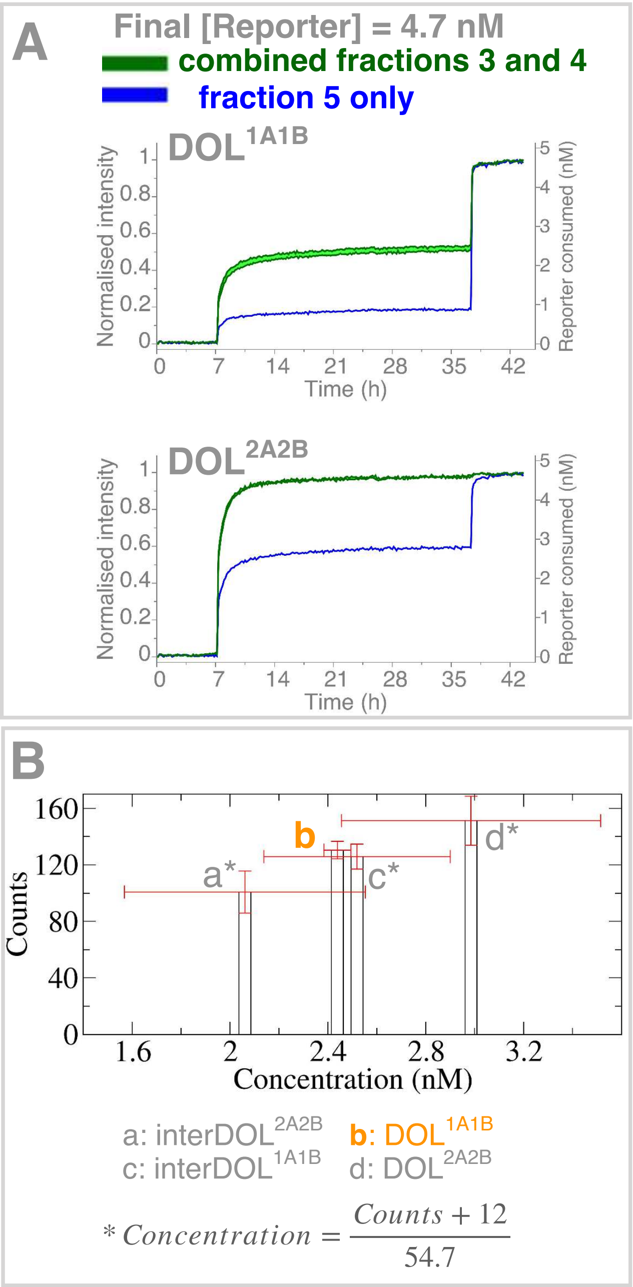


**Figure S4**: *Fluorescence for intra-DOL cases and concentration calculation*. **(A)** Similar to plate reader experiment in Figure 2D, kinetic curves are shown and compared in each case for pooled 3+4 fraction vs. fraction 5. **(B)** Counts vs. concentration bar plot shown for different DOL cases. $DOL^{1A1B}$ TEM data was used to obtain a general relationship between counts and concentration which was then used to predict concentrations for cases marked with * from their respective TEM data. More details in Supplementary Section 4.

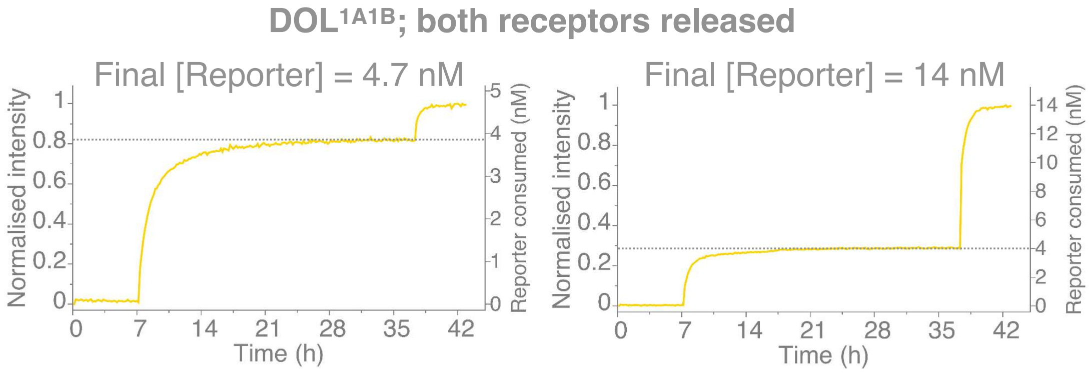


**Figure S5**: *Excess reporter complex does not block receptors*. Same DOL sample was reacted with two different reporter complex concentrations. In both the cases the saturation end points are marked with dotted horizontal lines. See related discussion in Supplementary Section 3.

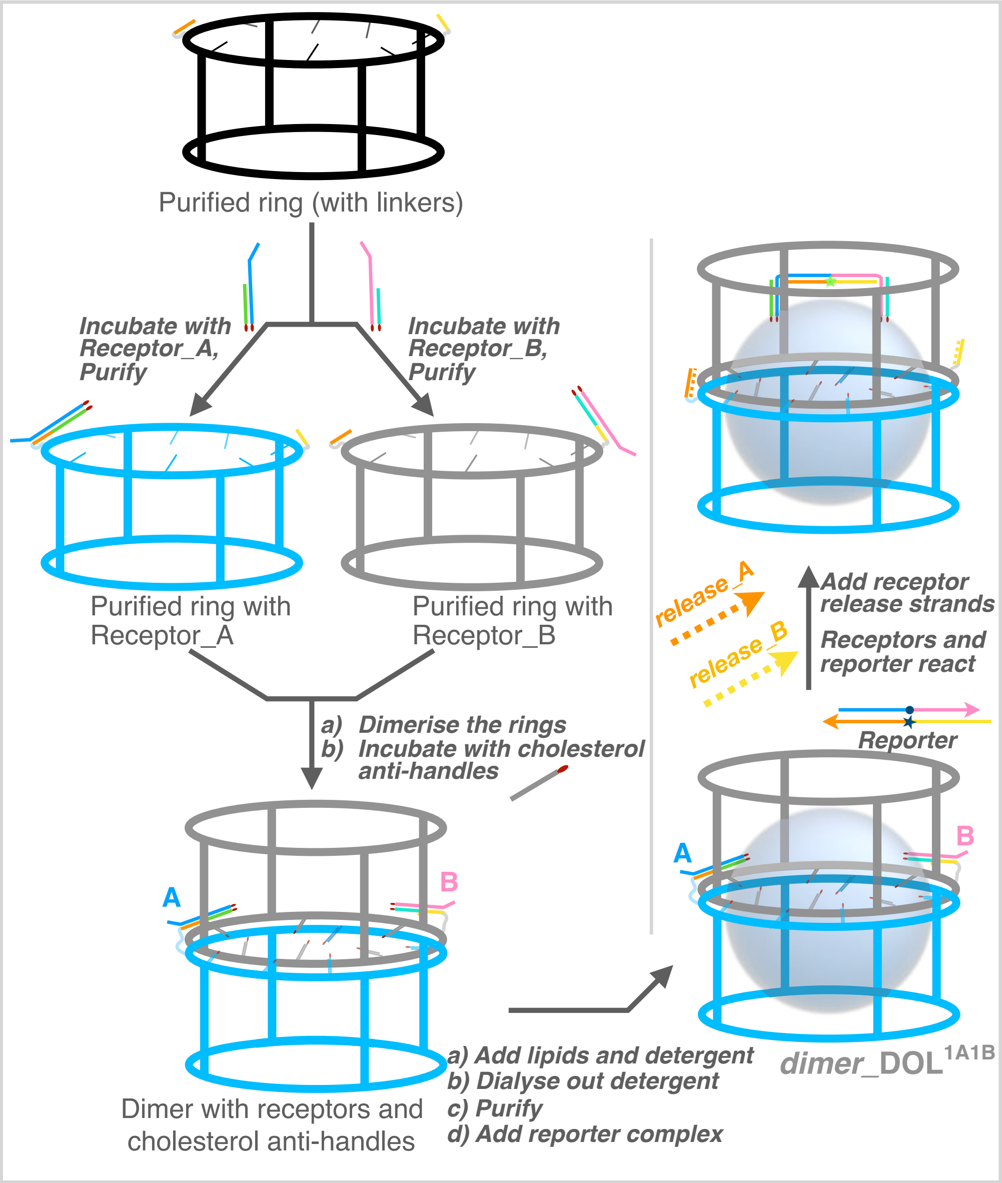


**Figure S6**: *Dimer ring platform*. Initially each ring was tethered with only one type of receptor and purified. Both the rings were later dimerised with the help of different (thus orientation is maintained) complementary staple extensions from one ring to the scaffold of the other ring. Overall the strategy remains similar to as explained in Figure 1.

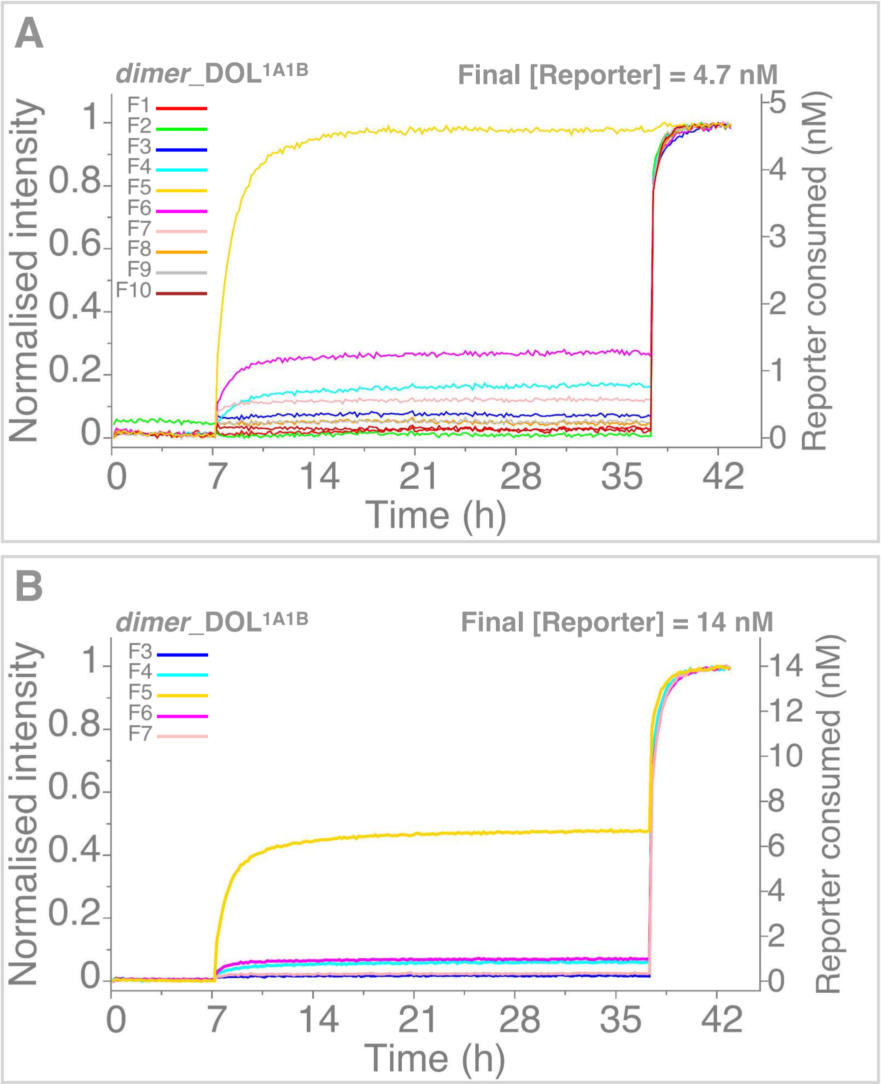


**Figure S7**: Analysing fractions obtained after isopycinic purification for dimer_DOL$^{1A1B}$ case.

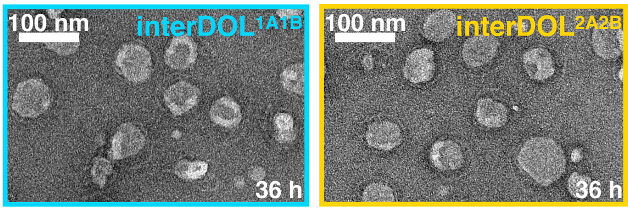


**Figure S8**: TEM images for inter-DOL cases. Samples were taken after the completion of the plate reader experiment shown in Figure 2E.

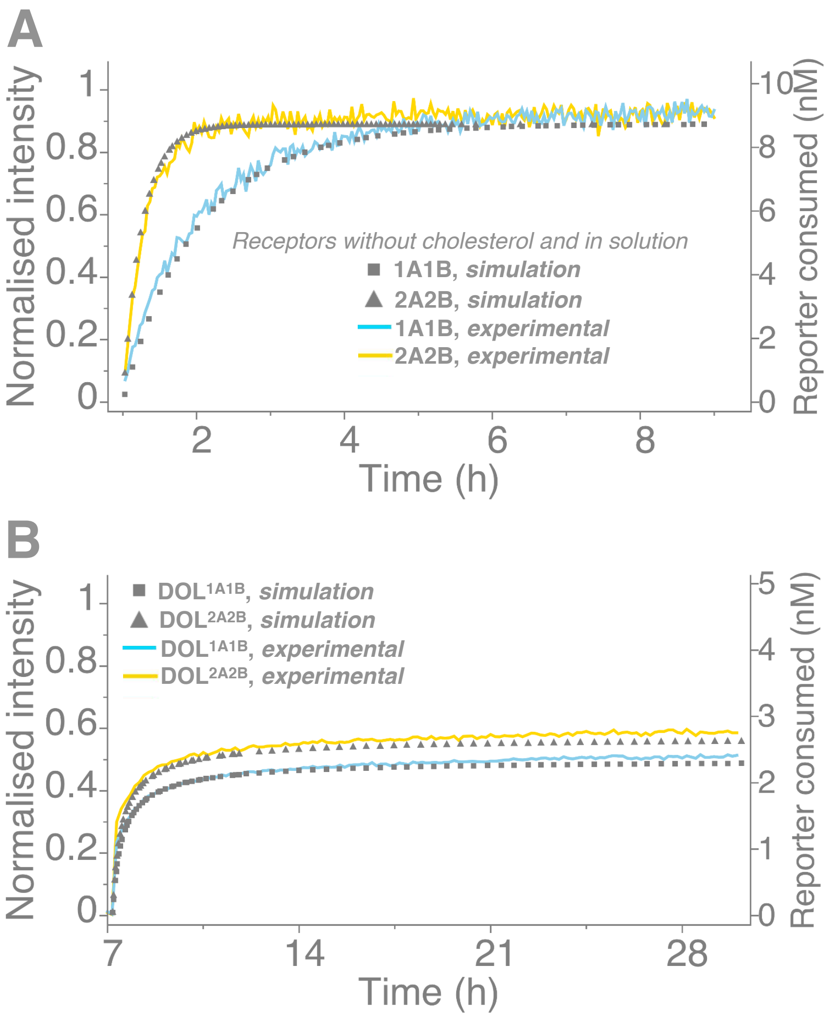


**Figure S9**: *Simulation curves*. **(A)** Experimental fluorescence kinetics curves in the solution case (Figure 2F) are plotted along with the fitted curves derived from the model discussed in Supplementary Section 6. For curve fitting data points from 1-9 h were used. **(B)** Similar to A, experimental (Figure 2D) and fitted curves shown for intra-DOL cases using data points 7-30 h.
